# Structure of an ant-myrmecophile-microbe community

**DOI:** 10.1101/2021.10.04.462948

**Authors:** Elena K. Perry, Stefanos Siozios, Gregory D. D. Hurst, Joseph Parker

## Abstract

Superorganismal ant colonies play host to a menagerie of symbiotic arthropods, termed myrmecophiles, which exhibit varying degrees of social integration into colony life. Such systems permit examination of how animal community interactions influence microbial assemblages. Here, we present an ecologically and phylogenetically comprehensive characterization of an ant-myrmecophile-microbe community in Southern California. Using 16S rRNA profiling, we find that microbiotas of the velvety tree ant (*Liometopum occidentale*) and its cohort of myrmecophiles are distinguished by species-specific characteristics but nevertheless bear signatures of their behavioral interactions. We found that the host ant microbiome was diverse at all taxonomic levels; that of a myrmecophilous cricket was moderately diverse, while microbiotas of three myrmecophilous rove beetles (Staphylinidae), which have convergently evolved symbiosis with *Liometopum*, were dominated by intracellular endosymbionts. Yet, despite these compositional differences, similarities between ant and myrmecophile microbiotas correlated with the nature and intimacy of their behavioral relationships. Physical interactions such as grooming and trophallaxis likely facilitate cross-species extracellular microbial sharing. Further, phylogenetic comparisons of microbiotas from myrmecophile rove beetles and outgroups revealed a lack of co-cladogenesis of beetles and intracellular endosymbionts, and limited evidence for convergence among the myrmecophiles’ intracellular microbiotas. Comparative genomic analyses of the dominant *Rickettsia* endosymbiont of the most highly socially integrated myrmecophile imply possible functions unrelated to nutrient-provisioning in the host beetle’s specialized lifestyle. Our findings indicate that myrmecophile microbiotas evolve largely independently of the constraints of deep evolutionary history, and that the transition to life inside colonies, including social interactions with hosts, plays a significant role in structuring bacterial assemblages of these symbiotic insects.

## Introduction

Insects constitute the bulk of global animal biodiversity and form integral components of both terrestrial and freshwater ecosystems (Scudder, 2017). Although many factors have been proposed to underlie insect diversification (Grimaldi & Engel, 2005), studies in numerous taxa have revealed that partnerships with symbiotic bacteria have been key to facilitating the exploitation of novel habitats and trophic resources (Engel & Moran, 2013). Insect-associated microbes have variously been shown to enable host survival on recalcitrant food sources (Douglas, 1998; Moran, McCutcheon, & Nakabachi, 2008; Russell et al., 2009; Salem et al., 2017; Zientz, Dandekar, & Gross, 2004), to synthesize chemical defenses (Piel, 2002), to influence communication, mating behavior and reproduction (Engl & Kaltenpoth, 2018; Wada-Katsumata et al., 2015), and to confer protection against parasites (Kaltenpoth & Engl, 2014; Koch & Schmid-Hempel, 2011; Oliver, Russell, Moran, & Hunter, 2003). Yet, despite efforts to characterize the many adaptive (and non-adaptive) roles that symbiotic microbe communities play in insect biology, knowledge of the converse relationship—how host ecology shapes the assembly and composition of the microbiome—remains scarce.

Both comparative studies, as well as experiments involving a small number of model insect species, indicate that host microbiotas can be influenced by habitat (Park et al., 2019; Yun et al., 2014), diet (Colman, Toolson, & Takacs-Vesbach, 2012; Majumder et al., 2020; Mason et al., 2020; Renelies-Hamilton, Germer, Sillam-Dussès, Bodawatta, & Poulsen, 2021), developmental stage (Jennings, Korthauer, Hamilton, & Benoit, 2019; Yun et al., 2014), and evolutionary history (Brooks, Kohl, Brucker, Opstal, & Bordenstein, 2016; R. T. Jones, Sanchez, & Fierer, 2013). Such studies have typically focused either on a single insect taxon, or have treated multiple insect taxa as separate, non-interacting entities. Yet, many of the central roles that insects play within the biosphere involve their interactions with other animal species. Such relationships—from predator-prey interactions to parasitic and mutualistic symbioses—are pervasive, and typify the biologies of many speciose insect clades (Bologna & Pinto, 2001; Feener & Brown, 1997; Godfray, 1994; Hölldobler & Wilson, 1990; Kathirithamby, 2009; Kistner, 1979, 1982; Kovarik & Caterino, 2005; Parker, 2016; Pierce et al., 2002; Stadler & Dixon, 2005). The nature of these interactions has in many cases evolved to become obligate, leading to extreme specialization of one insect species on a single or small number of partners (Beeren et al., 2021; Elmes, Barr, & Thomas, 1999; Hawkins, 1994; Komatsu, Maruyama, & Itino, 2009; López-Estrada, Sanmartín, Uribe, Abalde, & García-París, 2021; Maruyama & Parker, 2017; Strand & Obrycki, 1996). How such interspecies dependencies shape the evolution and composition of insect microbiotas remains an open question.

In other animal clades including mammals, growing evidence indicates that, in addition to host diet and phylogeny (Amato et al., 2019; Groussin et al., 2017), direct social interactions both within and between species can influence host-associated microbial communities. In chimpanzees, seasonal increases in social interaction correlate with increased similarity of gut microbiota (Moeller et al., 2016). In baboons, patterns of gut microbiome similarity can be predicted from social networks based on grooming interactions (Tung et al., 2015). Studies in humans have likewise correlated close social relationships and cohabitation with greater similarity of gut or skin microbiota (Dill-McFarland et al., 2019; Song et al., 2013). Similarly, dog owners share more members of their skin microbiota with their own pets than with other dogs, indicating that contact-mediated microbial exchange can occur between species, strongly influencing microbial community structure (Song et al., 2013). These observations imply that the degree of behavioral intimacy between interacting individuals—either within social groups or between species—may be a key parameter shaping animal microbiotas.

Here, we ask how the evolution of social behavioral relationships between insect species impacts their symbiotic bacterial communities. To address this question, we exploit a novel system: an ant species that is targeted by a cohort of socially parasitic insects. Many ants play keystone roles in terrestrial ecosystems and engage in relationships with diverse other arthropods (Hölldobler & Wilson, 1990; Parker & Kronauer, 2021). One pervasive mode of interaction occurs within the ant colony itself—a sheltered microhabitat that houses a concentration of resources in the form of ant brood and harvested food. Although ruthlessly policed against intruders, colonies are vulnerable to exploitation by an array of specialized arthropods that have evolved ways to evade recognition and gain entry (Kistner, 1979, 1982; Parker, 2016). Such organisms, termed “myrmecophiles”, are often obligately dependent on a single host ant species (Beeren et al., 2021; Elmes et al., 1999; Komatsu et al., 2009; Maruyama & Parker, 2017). To infiltrate colonies, myrmecophiles commonly employ deceptive strategies that permit them to forge intimate relationships with their unknowing hosts. Social integration of myrmecophiles typically hinges on chemical and behavioral adaptations, including mimicry of host ant pheromones (cuticular hydrocarbons; CHCs) (Akino, 2002; Bagnères, Blomquist, Bagnères, & Lorenzi, 2010; Beeren et al., 2018; Beeren, Schulz, Hashim, & Witte, 2011; Lenoir et al., 2012; Maruyama, Akino, Hashim, & Komatsu, 2009; Meer & Wojcik, 1982; Parker, 2016), or the secretion of so-called “appeasement compounds” that attenuate ant aggression and foster the myrmecophile’s acceptance into the nest (Akre & Hill, 1973; Cammaerts, 1992; Hölldobler, 1967, 1970; Hölldobler & Kwapich, 2019; Jordan, 1913; Parker & Grimaldi, 2014; Stoeffler, Tolasch, & Steidle, 2011). Once integrated into colonies, myrmecophiles can engage in intimate physical interactions with hosts, including reciprocal grooming, phoresy, and mouth-to-mouth feeding (trophallaxis) (Akre & Hill, 1973; Hölldobler, 1971; Hölldobler & Wilson, 1990; Kistner, 1979, 1982; Leschen, 1991; Parker, 2016). The obligate nature and extreme closeness of some ant-myrmecophile relationships provides a paradigm to explore how interspecies relationships influence microbial community structure in both the host and symbiont insects.

Our model ant-myrmecophile network centers on the ecologically dominant native ant of Southern California: the velvety tree ant, *Liometopum occidentale* (Formicidae: Dolichoderinae) (**Fig. 1A**). *Liometopum* ants form huge colonies numbering over one million workers, and patrol sectors of low- to mid-elevation oak and pine forest floor that can span tens to hundreds of meters in diameter (Hoey-Chamberlain, Rust, & Klotz, 2013; Wang, Patel, Vu, & Nonacs, 2010). Colonies of this ant house a large menagerie of myrmecophiles with different socially parasitic lifestyles and integrating strategies. By virtue of their distinct behaviors, these species permit insight into how different modes of interspecies interaction can impact symbiotic microbial assemblages. Prominent within *Liometopum* colonies are multiple species of rove beetle (Staphylinidae) that are obligately associated with this ant, and for which detailed understanding of ethology and chemical ecology has been obtained (Danoff-Burg, 1996). Crucially, each rove beetle species has independently evolved to socially parasitize *Liometopum* colonies: the species all belong to the same rove beetle subfamily, Aleocharinae (Staphylinidae), but have evolved into myrmecophiles from phylogenetically distant free-living ancestors belonging to different taxonomic tribes. This property permits comparisons of microbiota between the convergent myrmecophiles and outgroup lineages, potentially illuminating how the evolution of behavioral symbioses shapes the microbiome.

**Figure 1.**
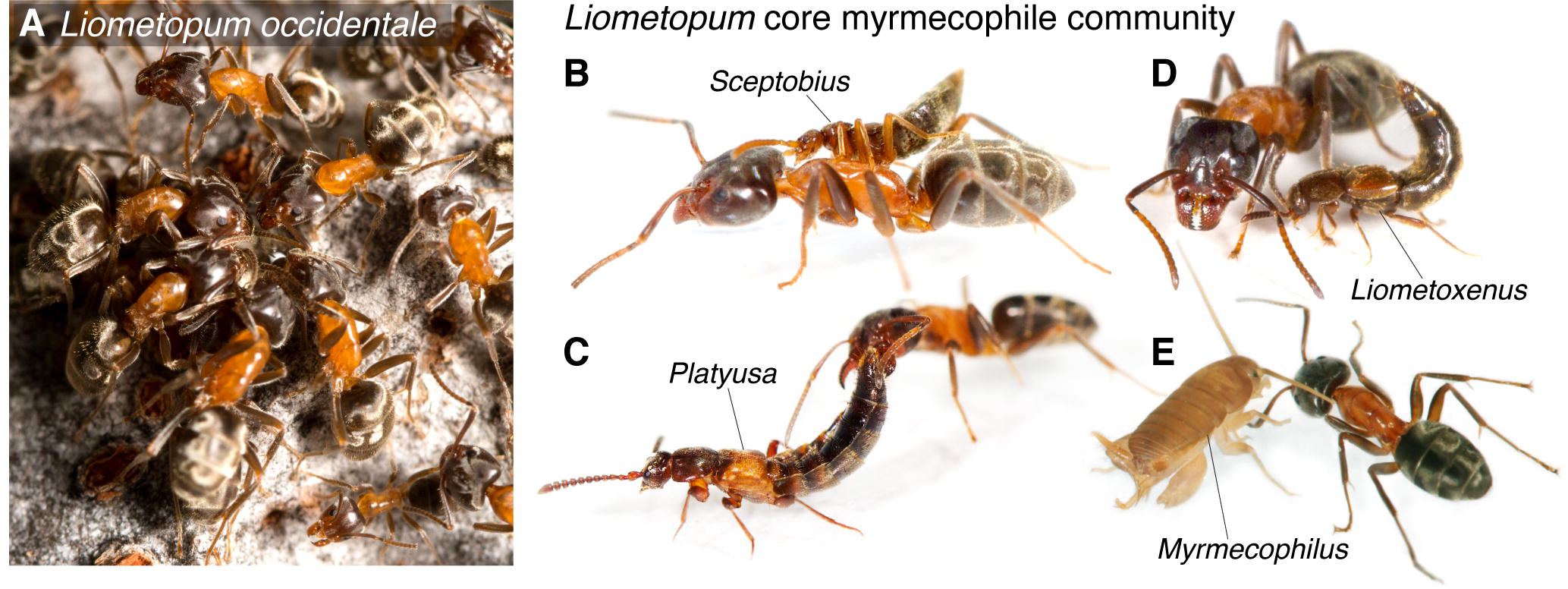
The *Liometopum* myrmecophile community. **A:** Workers of the velvety tree ant (*Liometopum occidentale*) foraging on a tree trunk. Credit: Kim Moore. **B:** *Sceptobius lativentris* rove beetle mounted on a *Liometopum* worker, performing grooming behavior. **C:** *Platyusa sonomae* rove beetle exuding appeasement secretion from abdominal gland to a *Liometopum* worker’s mouthparts. **D:** *Liometoxenus newtonarum* rove beetle with *Liometopum* nearby. **E:** *Myrmecophilus cf. manni* cricket interacting with a *Liometopum* worker.

In this study, we harness these attributes of the *Liometopum*-myrmecophile network to address how the social behaviors, evolutionary histories and microhabitats of interacting insect species contribute to structuring their symbiotic microbiotas. By extensive community sampling of core members of the myrmecophile network across colonies and localities, we have constructed a comprehensive, quantitative picture of the microbiome of an ant colony and its main social parasite symbionts. Incorporating knowledge of both myrmecophile behavior and microbiome data from outgroup relatives of key members of the myrmecophile network, we present evidence that the intimacy of interspecies social relationships is a major determinant shaping the evolution of insect bacterial communities.

## Materials and Methods

### Sample collection

Specimens of *Liometopum*, *Sceptobius*, *Myrmecophilus*, *Platyusa*, and *Liometoxenus* were collected in Southern California in May and July, 2019 (see “Collection localities” below). The specimens were collected into sterile 50 mL conical tubes, with each species in separate tubes, and stored at 4°C until processing later the same day. For dissected samples, specimens were transferred to sterile 1.5 mL microcentrifuge tubes and washed by vortexing at high speed for 10 s in 527 µL of sterile TE buffer (10 mM Tris-HCl, 1 mM EDTA, pH 8) just prior to dissection; the TE buffer fractions were retained as “body wash” samples and stored at -80°C. Whole body specimens were washed in the same manner (except for *Pella*, *Lasius*, *Drusilla*, and *Lissagria*), transferred to fresh sterile microcentrifuge tubes, and stored at -80°C. *Sceptobius* and *Platyusa* were dissected to separate the head and the gut from the rest of the body, while for *Liometopum* only the gut was separated from the rest of the body. For *Sceptobius*, due their small body size, three individuals were pooled per dissected body part sample (single individuals were used in the case of whole body samples) to ensure sufficient DNA yields due to the small size of the beetles. The dissections were performed in sterile Petri dishes using forceps and dissecting scissors that were sterilized in between samples with 10% bleach and flaming with ethanol. Dissected body parts were placed in fresh sterile 1.5 mL microcentrifuge tubes and immediately stored at -80°C until DNA extraction.

### Collection localities

Post Tree: USA, California, Altadena, Chaney Trail, 34.216293 -118.145922, 26 vii 2019, coll. E. Perry and J. Parker

First Tree: USA, California, Altadena, Chaney Trail, 34.216436 -118.148452, 26 vii 2019, coll. E. Perry and J. Parker

Rocks Tree: USA, California, Altadena, Chaney Trail, 34.217124 -118.154182, 26 vii 2019, coll. E. Perry and J. Parker

Sleepy Tree: USA, California, Altadena, Chaney Trail, 34.217456 -118.154426, 26 vii 2019, coll. E. Perry and J. Parker

Mile High Tree: USA, California, Altadena, Rubio Canyon Trail, 34.2053886, -118.1176927, 22 v 2019, coll. E. Perry and J. Parker

### DNA extraction and sequencing

DNA extractions were performed in batches of 11 samples, with one blank extraction per batch. To minimize the possibility of confounding batch effects with true differences between samples, each batch consisted of a random assortment of different sample types. To maximize bacterial DNA yield, the extractions were performed using a multistage protocol that involved grinding samples in liquid nitrogen, bead beating, and several rounds of phenol-chloroform extraction as described below.

Prior to starting extractions, 0.1 mm glass beads were sterilized by baking at 280°C for 4 hours. Working in a biological safety cabinet to maintain sterility, approximately 100 µL of beads were then transferred into each sample tube using a spatula sterilized with 50% bleach. After adding the beads, the samples (except for body wash samples) were ground to a fine powder in liquid nitrogen using blunt metal forceps that were sterilized in between samples with 50% bleach and flaming with ethanol. 527 µL of TE buffer was added to each tube immediately after grinding, except for body wash samples, which already consisted of the same volume of TE buffer; the latter were thawed on ice. Subsequently, 60 µL of 10% sodium dodecyl sulfate (SDS), 7.5 µL of proteinase K (20 mg/mL), and 6 µL of lysozyme (100 mg/mL) were added to each sample. The tubes were subjected to bead beating in a Disruptor Genie (Scientific Industries, Inc.) for 1 min at 4°C, and then incubated overnight at 37°C with shaking at 250 rpm.

The following day, 100 µL of 5 M NaCl was added and mixed thoroughly, followed by the addition of 80 µL of CTAB/NaCL solution (10% cetyltrimethylammonium bromide, 0.7 M NaCl) preheated to 65°C. The samples were incubated at 65°C for 10 min with periodic mixing by inversion. Next, 650 µL of 25:24:1 phenol/chloroform/isoamyl alcohol (pH 8) was added and mixed thoroughly by inversion, and the samples were spun at 13,000 rpm for 5 min in a microcentrifuge. The aqueous supernatant was transferred to a fresh 1.5 mL microcentrifuge tube and the extraction was repeated with 700 µL of 24:1 chloroform/isoamyl alcohol. Finally, the aqueous layer was extracted a third time with 500 µL of 24:1 chloroform/isoamyl alcohol. In addition, 650 µL of fresh TE buffer was added to the first round of phenol/chloroform/isoamyl alcohol tubes and the entire three-step extraction process was repeated, such that each organic fraction was extracted twice. Both aqueous fractions were then combined for each sample in a 1.5 mL DNA LoBind microcentrifuge tube (Eppendorf). DNA was precipitated overnight at -20°C with 0.6 vol isopropanol and 1 µL GenElute linear polyacrylamide (MilliporeSigma). Following precipitation, the DNA was pelleted by centrifuging at 13,000 rpm for 30 min at 4°C. The pellets were washed twice with 70% ethanol, then air dried and resuspended in 25 µL of 10 mM Tris buffer (pH 8). Resuspension was allowed to proceed overnight at 4°C prior to quantification of DNA yield using the Qubit dsDNA HS assay kit (ThermoFisher Scientific).

16S V4 amplification and Illumina library preparation were performed by Laragen, Inc., following the protocol recommended by the Earth Microbiome Project (Ul-Hasan et al., 2019). For approximately half of the samples (not limited to particular sample types), PCR amplification initially failed. Most of these samples ultimately yielded a detectable gel electrophoresis band upon the inclusion of 100 mg/mL bovine serum albumin in the reaction, or after further sample purification using the DNeasy PowerClean CleanUp kit (QIAGEN) and/or use of the KAPA3G Plant PCR kit (Roche) instead of Platinum Hot Start PCR Master Mix (ThermoFisher Scientific). The full list of sample treatments (excluding blank samples) is provided in **Table S1**. Amplified samples were sequenced on the Illumina MiSeq platform using the MiSeq v2 300 cycles kit (Illumina).

### 16S rRNA gene amplicon sequence processing and curation

MiSeq data were processed using the dada2 R package (version 1.14.1) (Callahan et al., 2016) to perform quality control (trimming and filtering) on sequences, infer exact amplicon sequence variants (ASVs), remove chimeras, and assign taxonomy to ASVs. The sequences were processed according to the dada2 pipeline recommendations (https://benjjneb.github.io/dada2/tutorial.html), but with a maximum of only one expected error allowed per read, and the truncation length set to 140. Taxonomy was assigned in dada2 using the SILVA reference database (version 132) (Quast et al., 2013).

ASVs identified as chloroplasts, mitochondria, or eukaryotic were removed from the data set using QIIME 2 (version 2020.2) (Bolyen et al., 2019). Additional putative contaminant sequences were identified and removed using the decontam R package (version 1.6.0) (Davis, Proctor, Holmes, Relman, & Callahan, 2018) in conjunction with phyloseq (version 1.30.0) (McMurdie & Holmes, 2013). ASVs were considered likely contaminants if they were identified as such both by frequency in negative controls vs. true samples and by prevalence across negative controls vs. true samples (using the default *p-*value threshold of 0.1). The blank DNA extractions served as the negative controls. For the frequency metric, the number of reads from each sample was used as a proxy for DNA concentration.

Following the removal of putative contaminants, all further sequence processing was performed in QIIME 2. First, the reference sequences were aligned and a phylogenetic tree was generated using the command “align-to-tree-mafft-fasttree.” Then, samples were rarefied to 1000 reads; five samples were discarded due to falling below this threshold. Alpha diversity metrics (Simpson index and chao1 estimate) were calculated, and beta diversity distance matrices (based on Bray-Curtis dissimilarity, Jaccard distance, and weighted and unweighted UniFrac distances) were generated. Inspection of principal coordinates analysis plots generated in QIIME 2 for the different distance metrics revealed no obvious clustering of samples based on differential treatment during sample cleanup or PCR amplification. For insect body part and whole body samples, beta diversity analyses were performed both before and after filtering out putative intracellular endosymbiont ASVs, while for body wash samples, the analyses were performed before and after filtering out all ASVs that appeared in any nest sample. Finally, after filtering out ASVs with fewer than 10 reads total across all samples to reduce the number of zeroes in the feature table, ASVs that were differentially abundant across *Liometopum*, *Sceptobius*, *Myrmecophilus*, and *Platyusa* samples (excluding putative intracellular endosymbionts and body wash samples) were identified using the QIIME 2 DS-FDR (discrete false discovery rate control) plugin with Kruskal-Wallis tests (Jiang et al., 2017).

### Genome sequencing, assembly, and annotation

For Illumina sequencing we used a single *Sceptobius lativentris* female (USA: California, Altadena, Eaton Canyon Falls Trail, 34°11’38.1′N 118°06’11.8′W vi.2017, coll. J. Parker). Genomic DNA was isolated via phenol/chloroform method, and integrity was assessed with a Bioanalyzer. An Illumina paired-end library (150bp) was prepared using the Illumina TruSeq DNA kit and sequenced on the Illumina HiSeq X platform by Iridian Genomes/J. Parker and collaborators. Reads were deposited on NCBI’s Sequence Read Archive (accession: SRR5909496). For long reads, total genomic DNA was extracted from 20 *Sceptobius* males and sequenced on an Oxford Nanopore minION flow cell at the Millard and Muriel Jacobs Genetics and Genomics Laboratory at Caltech. An initial assembly of the Illumina reads from the SRR5909496 library was assembled *de novo* using megahit v1.2.9 (D. Li, Liu, Luo, Sadakane, & Lam, 2015) and the contigs were binned using Metabat2 (Kang et al., 2019) based on tetranucleotide frequencies. A *Rickettsia* like bin was identified using CheckM v1.0.13 (Parks, Imelfort, Skennerton, Hugenholtz, & Tyson, 2015) and was used as a reference to map both short and long reads using minimap2 (H. Li, 2018). The *Rickettsia* like reads were extracted with samtools v1.9 (H. Li et al., 2009) and a hybrid assembly of Nanopore and Illumina reads was performed using the Unicycler pipeline v0.4.8 under the normal mode (Wick, Judd, Gorrie, & Holt, 2017). The raw reads (long and short) were mapped back to the assembly and the assembly was visually inspected for misassemblies. Genome annotation was performed with prokka v1.14.0 (Seemann, 2014) and Pfam domains were predicted with InterProScan v86.0 (P. Jones et al., 2014). Genomic contigs as well as the complete annotated genome and annotation is provided in **Files S1**, **S2** and **Table S5**.

### Phylogenomic analysis

The phylogenetic position of the *Rickettsia* endosymbiont of *Sceptobius lativentris* was estimated in relation to 86 publicly available genomes from all major *Rickettsia* groups based on the concatenated analysis of 267 single copy core proteins determined using OrthoFinder v2.3.11 (Emms & Kelly, 2019). Phylogenetic relationships were inferred using maximum likelihood as implemented in IQ-TREE 2.0.3 (Minh et al., 2020) under the JTT+F+R5 model which was selected using ModelFinder (Kalyaanamoorthy, Minh, Wong, Haeseler, & Jermiin, 2017). The robustness of the tree was finally assessed based on 1000 ultrafast bootstrap replicates (Hoang, Chernomor, Haeseler, Minh, & Vinh, 2017). The phylogenetic tree was visualized and annotated using iTOL (Letunic & Bork, 2007).

### Comparative genomic and metabolic analysis

A comparative genomic analysis across members of the Transitional group of *Rickettsia* was performed using anvi’o package v7 (Eren et al., 2021). The following parameters were used to create the pangenome of the transitional group: “--use-ncbi-blast --mcl-inflation 1.5”. The metabolic potential of RiSlat genome was estimated using the “anvi-estimate-metabolism” program from anvi’o package by computing the completion ratio of the predicted KEGG functional categories and compared to the other members of the transitional group. Heatmaps were prepared using the R package pheatmap (Kolde & Kolde, 2015). We computed the fraction of interrupted genes (genes which length was shorter than 80% of their best hit in the Swiss-Prot database) using the ideel method (https://github.com/mw55309/ideel). Finally, repeat density (repeats ≥ 1500 bp and at least 95% identity) in RiSlat genome and the synteny between RiSlat genome and complete reference genomes from the transitional group of *Rickettsia* was computed and visualized using MUMmer v3 (Kurtz et al., 2004).

### Data analysis and statistics

Non-metric multidimensional scaling (NMDS) ordinations were generated using the R package vegan (version 2.5-6) (Oksanen et al., 2015). Permutational multivariate analysis of variance (PERMANOVA) was performed on distance matrices using the “adonis” function in vegan with 999 permutations. Distributions of distances between different sample types were compared in R either using the Wilcoxon test followed by the Benjamini-Hochberg method to control the false discovery rate for multiple comparisons, or using the Kruskal-Wallis test followed by Dunn’s test for pairwise comparisons. Dunn’s test was performed using the function “dunnTest” from the package FSA (Ogle, Wheeler, & Dinno, 2020). Complete information on the statistical tests used in each figure is provided in **Table S2**. Plots in **Figs. 2-4** and **Figs. S1-S3** were generated in R with the package ggplot2 (version 3.3.1) (Wickham, 2011) except that the package pheatmap (Kolde & Kolde, 2015) was used to generate the heatmap in **Fig. 3C**, and the package bipartite (version 2.15) (Dormann, Gruber, & Fründ, 2008) was used to generate the bipartite network plot in **Fig. 4B**.

**Figure 2.**
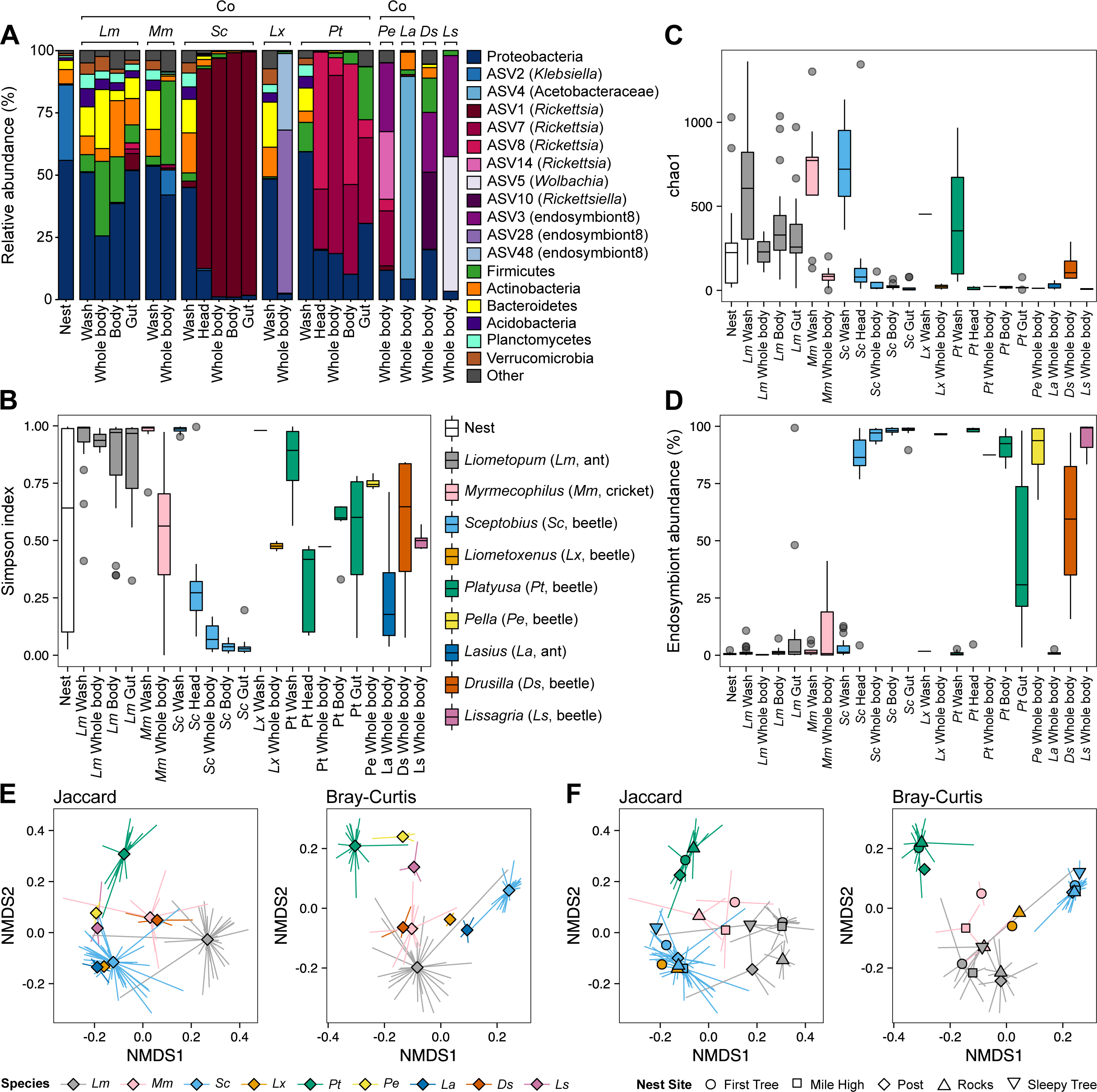
Microbial diversity of *Liometopum* and core myrmecophiles. **A:** Bar plots showing relative abundance of different bacterial taxa in the studied species. Samples belonging to the same insect species and sample type were combined additively. *Lm* = *Liometopum* (ant), *Mm* = *Myrmecophilus* (cricket), *Sc* = *Sceptobius* (beetle), *Lx* = *Liometoxenus* (beetle), *Pt* = *Platyusa* (beetle), *Pe* = *Pella* (beetle), *La* = *Lasius* (ant), *Ds* = *Drusilla* (beetle), *Ls* = *Lissagria* (beetle). “Co” indicates species that cohabitate. Taxa bar plots for individual samples are provided in **Fig. S1. B-D:** Distributions of alpha diversity metrics for each sample type, as quantified by the Simpson index (**B**), chao1 species richness estimate (**C**), and relative abundance of putative endosymbionts (**D**). The color legend in panel B also applies to panels C and D. **E-F:** NMDS ordination of the insect samples (not including wash samples) based on either Jaccard distance (unweighted by abundance) or Bray-Curtis dissimilarity (weighted by abundance). Diamonds (E) or other shapes (F) mark the centroids of each species or sample group, while line segments connect the centroids to the individual data points. Centroids and line segments are colored by the species of origin in both E and F, and in F the marker shape for the centroid of each group denotes the nest site where the samples were collected. Not all samples from E are shown in F, as site information was only available for *Liometopum* and its associated myrmecophiles.

**Figure 3.**
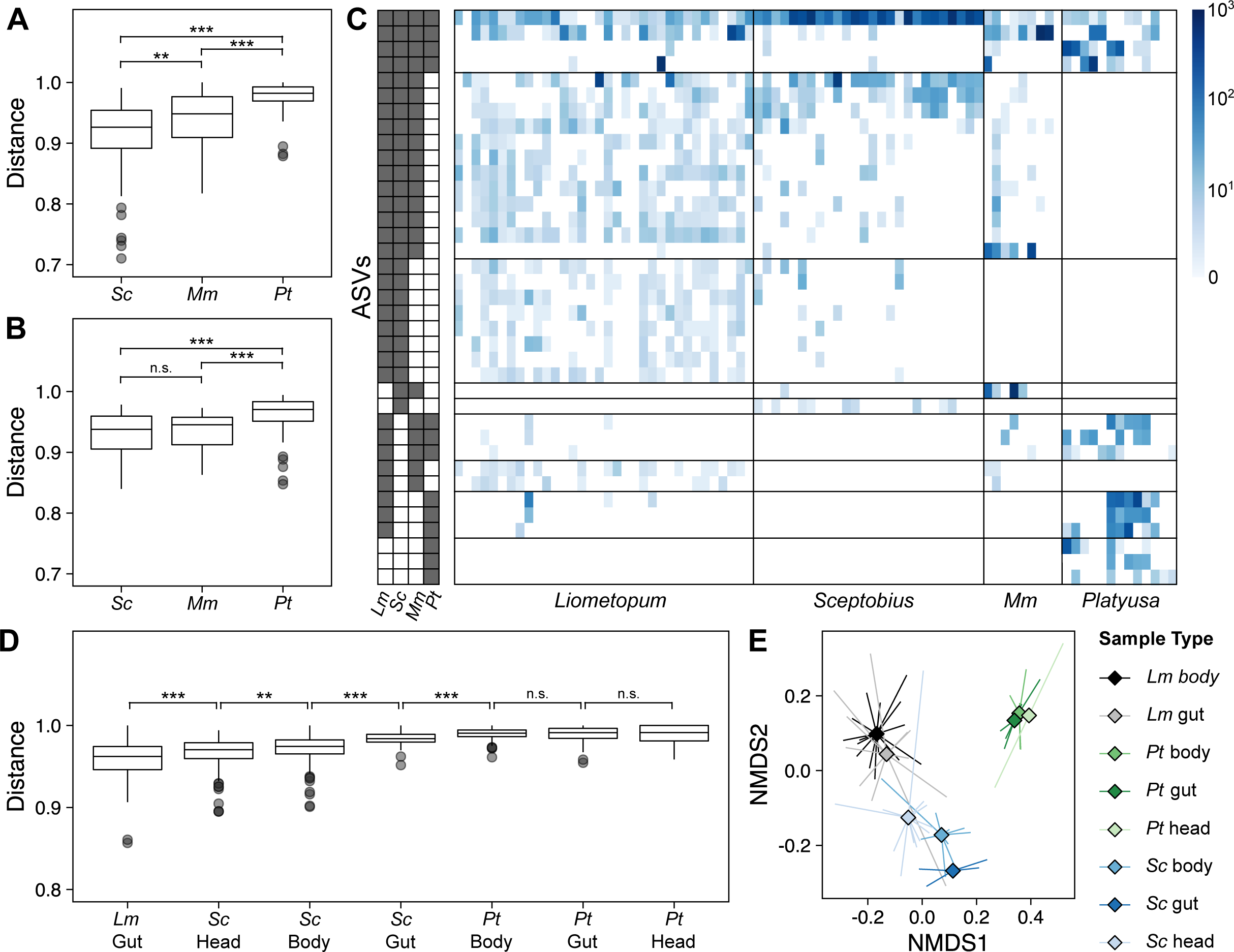
Social interactions explain microbial sharing between ants and myrmecophiles. **A:** Jaccard pairwise distances for *Sceptobius* (*Sc*), *Platyusa* (*Pt*), and *Myrmecophilus* (*Mm*) versus *Liometopum,* within nest sites, without endosymbionts. All sample types except wash samples (i.e. whole body samples in addition to dissected body parts where applicable) were included for each species. **B:** Jaccard pairwise distances for body washes of *Sceptobius* (*Scep*), *Platyusa* (*Plat*), and *Myrmecophilus* (Cric) versus *Liometopum*, within nest sites, after removing all ASVs that appeared in any Nest sample. **C:** Visualization of ASVs that were differentially abundant across ant, *Sceptobius*, cricket, and *Platyusa* samples, as determined by Kruskal-Wallis tests using the DS-FDR method to control the false discovery rate to 0.1. Left: Visualization of the presence (gray)/absence (white) of each ASV in samples pooled by insect species. Each row represents an ASV and the columns from left to right represent ant, *Sceptobius*, cricket, and *Platyusa* respectively. Right: Heatmap showing the normalized read counts for each ASV in each sample. Each row represents an ASV (same order as on the left), each column represents an individual sample, and the heatmap cells are colored by read count. See **Table S3** for the ASV identities and statistical values. **D:** Jaccard pairwise distances between different myrmecophile body parts and ant bodies, without endosymbionts. **E:** NMDS of Jaccard pairwise distances for *Sceptobius* (*Sc*), *Platyusa* (*Pt*), and *Liometopum* (*Lm*) by body part, without endosymbionts. Diamonds represent the centroid for each sample type, and line segments connect the centroids to the individual data points.

**Figure 4.**
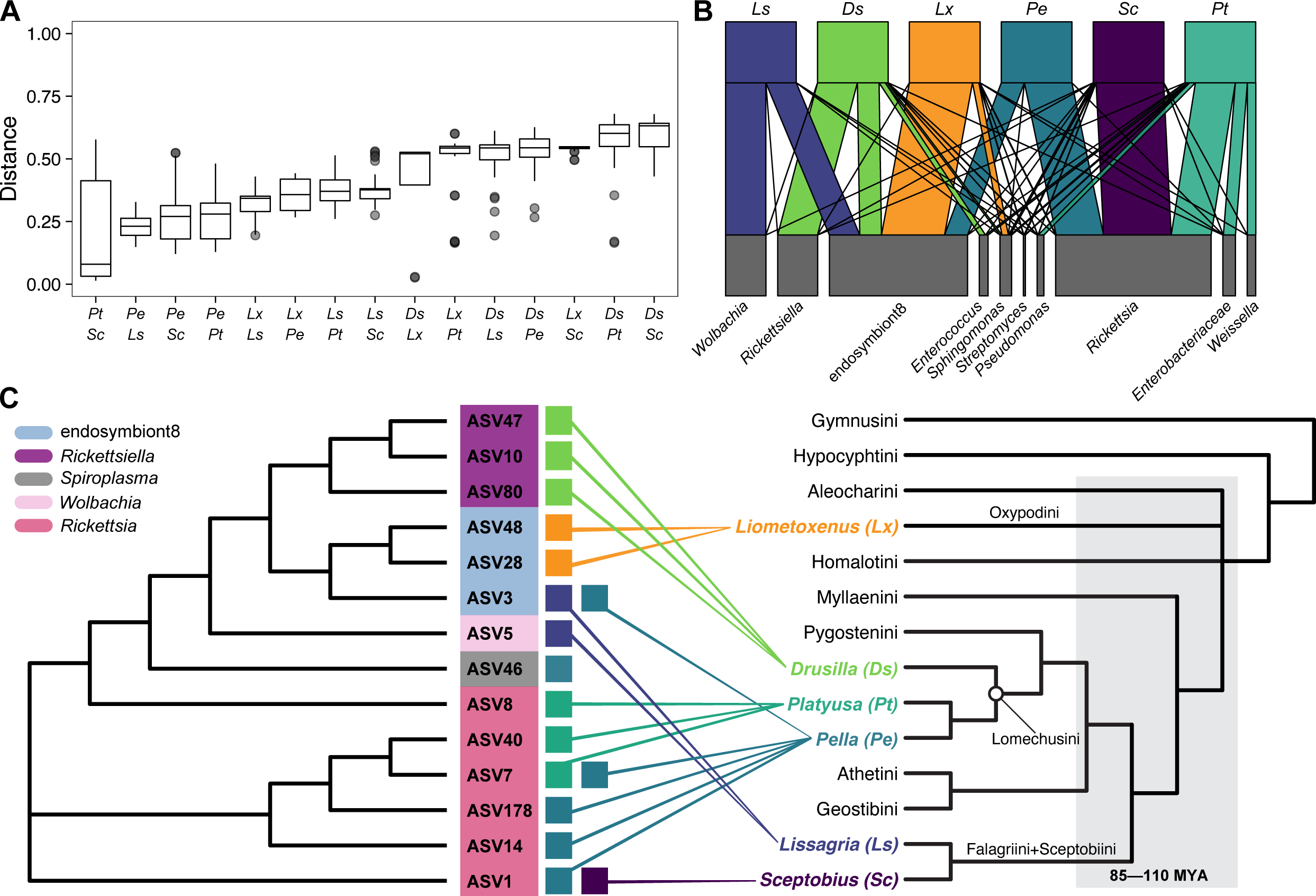
Relationship between myrmecophile phylogeny and microbiota composition. **A:** Distributions of weighted UniFrac distances between individuals of different species pairings within the staphylinid beetles in this study. All sample types were included except for wash samples. **B:** Bipartite network plot showing taxonomic overlap of the microbiotas belonging to the different staphylinid beetles. Only the top ten taxa by abundance across these samples are shown. Samples derived from the same species were pooled additively and the taxon abundances were normalized by the number of total reads for each beetle species. **C:** Comparison of the cladogram of the putative endosymbionts associated with the staphylinid beetles and the cladogram of the beetles used in this study together with other major tribes of Aleocharinae, showing a lack of codiversification of the putative endosymbionts with their beetle hosts. The period of extensive tribal-level cladogenesis, 85–110 MY ago, is shaded grey (tree based on Maruyama & Parker, 2017; Orlov, Newton, & Solodovnikov, 2021).

## Results

### The *Liometopum*-myrmecophile community

Our study sites are located in the foothills of the San Gabriel Mountains within Angeles National Forest, LA County, California, where *Liometopum occidentale* is the predominant ant species. We have found that colonies of this ant play host to a “core” community of four obligate myrmecophiles, which occur in and around most, if not all nests, and exhibit distinct behavioral interactions with host workers. Three members of the core community belong to the hyperdiverse rove beetle subfamily Aleocharinae (Coleoptera: Staphylinidae)—a clade of ∼17,000 species in which numerous lineages have evolved myrmecophilous lifestyles (Kistner, 1979; Maruyama & Parker, 2017; Parker, 2016; Seevers, 1965; Yamamoto, Maruyama, & Parker, 2016). One of these species, *Sceptobius lativentris* (Aleocharinae: Sceptobiini) ranks among the most socially integrated myrmecophiles known. *Sceptobius* beetles live in close physical intimacy with host ants, climbing onto workers’ bodies where they spend up to 80% of their adult lives (**Fig. 1B; Video S1**). Attaching to workers enables the beetle to groom the ant with its legs—an adaptive behavior that functions to transfer the ant’s CHCs onto the beetle’s own body. Ant grooming permits *Sceptobius* to achieve identical chemical resemblance to its host and gain social acceptance, reflected in the presence of beetles throughout the nest, including the brood galleries. *Sceptobius* is believed to feed on ant brood as well as via trophallaxis with workers, and may also perform “strigilation”—grazing on material attached to worker body surfaces (Danoff-Burg, 1996). The beetles are unable to survive for longer than ∼24 hours without direct, physical interactions with *Liometopum* workers (Danoff-Burg, 1996). *Sceptobius* thus embodies the phenotypic extreme of socially complex symbiotic lifestyles, and is the most tightly associated of the core *Liometopum* myrmecophiles.

The second myrmecophile, the aleocharine *Platyusa sonomae* (Aleocharinae: Lomechusini), is behaviorally distinct to *Sceptobius*: it occurs in the vicinity of colonies and along ant foraging trails but appears never to penetrate inside the nest. *Platyusa* interacts physically but transiently with ants: the beetles appease and pacify aggressive workers with attractive secretions from an abdominal gland (**Fig. 1C**). Additionally, physical interactions occur through direct predation on worker ants. Unlike *Sceptobius*, *Platyusa* can be cultured independently of its host ant in the laboratory and fed alterative diets. The third aleocharine, *Liometoxenus newtonarum* (Aleocharinae: Oxypodini) is found predominantly at the mouths of nests where ant density is high, but we have not observed this species deeper in the colony. *Liometoxenus* intermingles with ant workers (**Fig. 1D**), but does not groom them; instead, *Liometoxenus* produces a volatile, ester-rich cocktail from an abdominal gland that acts at a distance, ‘intoxicating’ ants and impeding their locomotion. Like *Platyusa*, *Liometoxenus* also predates on worker ants. Unlike the other core myrmecophiles which are found commonly year-round, *Liometoxenus* adults appear only during spring, and are somewhat rarer. The final core myrmecophile is the ant cricket, *Myrmecophilus cf. manni* (Orthoptera: Myrmecophilidae). Behavioral studies of this and other *Myrmecophilus* species, as well as our own observations, have shown that the crickets interact closely with ants (**Fig. 1E**), feeding via both trophallaxis with workers and also via opportunistic strigilation, wherein the cricket rapidly approaches the ant and briefly grazes its body surface before retreating (Hebard, 1920; Henderson & Akre, 1986; Morton, 1900). These actions parallel some behaviors of *Sceptobius*. However, *Myrmecophilus* is not phoretic upon *Liometopum* workers, does not groom them, and based on our observations, does not access the internal parts of colonies but instead inhabits nest entrances and trails.

The distinct trophic and physical interactions between the four myrmecophiles and their shared *Liometopum* host ant motivated us to ask how their contrasting lifestyles are reflected in their microbial symbionts. We generated bacterial 16S rRNA metabarcoding libraries from *Liometopum* workers and each of the core myrmecophile species, sampling specimens from five ant colonies in two field sites in the Angeles National Forest (see **Materials and Methods**). For all species, we sequenced whole insect bodies, as well as microbes washed from the body surface. Further, given the myrmecophiles’ diverse modes of social interaction and feeding behavior, we also sequenced specific microbial communities from the gut and body of *Liometopum* host workers, and from the gut, head and body of the myrmecophiles *Sceptobius* and *Platyusa*. The latter two species display the most divergent trophic behaviors, and in behavioral terms represent the most and least extreme degrees of social integration into the host colony. Since each of the rove beetles has convergently evolved to target *Liometopum*, we additionally asked to what extent the microbial communities of *Sceptobius* and *Platyusa* are influenced by their evolutionary histories. To do so, we sequenced whole-body microbiota from free-living outgroups of both species: for *Sceptobius,* we obtained the North American aleocharine *Lissagria laeviuscula* (tribe Falagriini: the sister tribe of Sceptobiini, or from which Sceptobiini is a derived lineage; Danoff-Burg, 1994, 2002; Maruyama & Parker, 2017), while for *Platyusa*, we obtained the European aleocharine *Drusilla canaliculata* (tribe Lomechusini). We also obtained specimens of the European aleocharine *Pella cognata* (tribe Lomechusini)—a myrmecophilous close relative of *Platyusa*, which is associated with a different host ant, the formicine *Lasius fuliginosus* (the microbiome of which we also sequenced). In total, we sequenced 230 libraries from the host ant, the four core myrmecophiles, environmental nest material from each sampled colony, blank (control) samples, as well as replicate libraries from each outgroup species and host ant.

### Microbiome of the velvety tree ant

We determined the taxonomic composition and alpha diversity of each species’ microbiota by quantifying bacterial 16S rRNA amplified sequence variants (ASVs)—a proxy for species (**Fig. 2A**). Of all species surveyed, the host ant, *Liometopum occidentale*, possessed the most diverse microbiota by two different metrics: the Simpson index, which is influenced by both species richness and evenness within each sample, and the chao1 estimate of total species richness (**Fig. 2B, C**). In addition to alpha diversity at the ASV level, the *Liometopum* microbiota was also the most diverse at the phylum level, encompassing Proteobacteria, Firmicutes, Actinobacteria, Bacteroides, Acidobacteria, Planctomycetes, and Verrucomicrobia—each comprising at least 1% of the 16S reads from pooled ant body or gut samples (**Fig. 2A**). Taxonomic composition at the phylum level was qualitatively similar across pooled *Liometopum* wash, body and gut samples, but generally distinct from the nest samples. For example, the majority of the latter contained a high proportion of reads from a single ASV classified as *Klebsiella*, which was nearly absent from the *Liometopum* samples, while many *Liometopum* samples had high proportions of Firmicutes, which were nearly absent from nest samples (**Fig. 2A**, **Fig. S1**). These distinctions suggest that *Liometopum*’s microbiota is typically distinct from its own nest material, and that physiological characteristics of *Liometopum* likely influence the assembly of its microbiota.

Despite coarse similarity among microbiota samples at the phylum level, however, *Liometopum* guts did not possess a distinctive “core” microbiota across samples, defined here as the set of ASVs present in at least 90% of the samples of a given species and body part (see **Table S1** for ASV abundances, both per sample and means per sample type). Across *Liometopum* body samples, only ASV6 (*Sphingomonas*) was highly prevalent, albeit at a low relative abundance (median of 0.55%). This same ASV was also present in several other species at a similar abundance (**Table S1**), and we cannot rule out this it is a contaminant (Salter et al., 2014). We also found no evidence for obligate, intracellular endosymbionts among the *Liometopum* microbiota. Such endosymbionts are typically known from ant species that utilize more specialized food sources than the generalized, omnivorous/scavenger diet of *L. occidentale* (Feldhaar et al., 2007; Gil et al., 2003).

### Species-specific characteristics of myrmecophile microbiotas

With the exception of body wash samples, microbial alpha diversities of the four core myrmecophiles were all markedly lower than that of the host ant. The cricket, *Myrmecophilus*, harbored the greatest diversity at both the ASV and phylum levels, with large proportions of both Proteobacteria and Firmicutes (42% and 34% respectively across pooled samples; **Fig. 2A**, **Fig. S1**). Median Simpson index and chao1 estimates from whole body samples were 0.56 and 83, respectively (**Fig. 2B, C**). Like its host ant, however, *Myrmecophilus* also did not possess a distinctive, core microbiota at the ASV level (**Table S1**). *Sceptobius* exhibited the least diverse microbiota, with Simpson index values of 0.03-0.27, depending on the body part, and correspondingly low chao1 estimates (**Fig. 2B, C**). The two remaining rove beetles, *Platyusa* and *Liometoxenus*, both exhibited intermediate Simpson index values of 0.48 (**Fig. 2B**). However, the chao1 species richness estimates for *Platyusa* and *Liometoxenus* were both low (16.5 and 22, respectively; excluding wash samples), indicating that these species’ higher Simpson index values likely reflected greater evenness of the microbial communities relative to *Sceptobius* rather than greater species richness.

The most striking feature of the microbiotas of all three staphylinids was the high proportions of reads from putative intracellular endosymbionts (**Fig. 2D**, **Fig. S1**). For example, >95% of 16s reads from *Sceptobius* bodies and guts belonged to a single variant of the intracellular Proteobacteria genus *Rickettsia*. This variant, ASV1, also accounted for 80% of reads from *Sceptobius* heads, indicating its likely presence throughout the body (**Fig. 2A**; **Table S1**). Similarly, *Rickettsia* species dominated the *Platyusa* microbiota, with three variants, ASV7, ASV8, and ASV40, that were differentially abundant across different body parts (**Fig. 2A**, **Fig. S1**; **Table S1**). ASV8 was more abundant in the head (median relative abundance of 66.8%) and body (43.1%) compared to the gut (2.9%), while ASV40 was most prevalent and abundant in *Platyusa* body samples (median relative abundance of 7.2%), but absent from head samples. ASV7 was generally present in similar proportions in all types of sequenced body parts (median relative abundance of 28-44%) and was also present at a comparable frequency in one undissected *Platyusa* whole body (**Fig. 2A**, **Fig. S1**; **Table S1**). Although *Liometoxenus* lacked *Rickettsia*, its microbiota was nevertheless dominated by two variants of a putative endosymbiont named “endosymbiont8” in the SILVA reference database (Quast et al., 2013). These ASVs, 28 and 48, belong to the Enterobacteriaceae family with 91-92% BLAST similarity to *Providencia* (the closest genus match), and collectively account for 95% of reads from this beetle species (**Fig. 2A, D**; **Table S1**). Beyond these dominant endosymbionts, the rove beetles possessed few other core microbial taxa: the aforementioned *Sphingomonas* (ASV6) was the only other consistently recovered non-intracellular bacterium in *Sceptobius,* specifically in the body and gut, and was present at low levels (0.2 and 0.5% median relative abundance respectively), while *Platyusa* did not possess any core ASVs other than ASV8 in either the head, body, or gut (**Table S1**). As for *Liometoxenus,* two *Arhodomonas* taxa, ASV6 and ASV39 were the only other two sequence variants present in both samples, at median relative abundances of 1.7 and 0.25% respectively (**Table S1**).

### Both species and colony identity impact host ant and myrmecophile microbiotas

NMDS ordinations based on both Jaccard distances and Bray-Curtis dissimilarities revealed that samples tended to cluster by species, with some overlap (**Fig. 2E**). PERMANOVA analysis revealed a statistically significant effect of species (*p* = 0.001; **Table S2**), confirming that the ant and its myrmecophiles possess distinguishable microbiotas. Given the relatively low shared diversity of microbial taxa across individuals within each insect species, however, we asked whether microbiotas were also potentially shaped by colony-level factors. Differences between source colonies may derive from ‘background’ environmental microbes that are not necessarily symbiotic. Alternatively, there may be colony-level differences in symbiotic microbiota, reflecting differential acquisition of microbes that are associated with host insect physiology. To distinguish between these possibilities, we performed NMDS ordination (**Fig. S2A**) and PERMANOVA on samples of collected nest material to determine whether background environmental microbes differed between colonies. For both Jaccard distance and Bray-Curtis dissimilarity, PERMANOVA revealed a statistically significant effect of nest site (*p* < 0.05; Table S2), with R^2^ ranging from 0.34 (Jaccard distance) to 0.42 (Bray-Curtis dissimilarity), reflected in moderate clustering in the ordination plots (**Fig. S2A**), confirming that environmental microbial communities indeed differed between colonies.

Before comparing insect samples from the different colonies, we verified whether the washes had effectively decreased reads from bacteria present in the nest environment, which we postulated would represent primarily transient, low-abundance members of the background microbiota rather than stably associated symbionts. Indeed, by both Jaccard distance and Bray-Curtis dissimilarity, wash samples were significantly more similar than insect body part samples to nest material samples (**Fig. S2B**). With the latter metric, there was also a long tail of samples with relatively low dissimilarity between insect body parts and nest material, possibly because unlike Jaccard distance, Bray-Curtis dissimilarity is weighted by abundance and therefore less sensitive to low-abundance community members (Birtel, Walser, Pichon, Bürgmann, & Matthews, 2015). Nevertheless, these results suggest that the washes successfully removed at least some environmentally derived bacteria. PERMANOVA analysis indicated that even after removing wash samples from the dataset, colony identity still had a significant effect (*p* = 0.001; **Table S2**) on the microbiota associated with the ants and myrmecophiles, based on Jaccard distances, although insect species explained a larger proportion of variation than nest site. These trends can be seen in the NMDS ordination of samples grouped by either insect species (**Fig. 2E**) or nest site (**Fig. 2F**). The statistically significant effect of colony identity was primarily driven by body samples, as PERMANOVA did not reveal a significant effect of location when the analysis was restricted to either gut or head samples. The PERMANOVA analysis revealed that the interaction between nest site and insect species was statistically significant *(p* = 0.008; **Fig. 2F**; **Table S2**), suggesting that inter-colony differences in the insect microbiotas were not driven by the same sets of microbes across the different species. These data uncover colony-to-colony variation in the symbiotic microbiota associated with each insect species—variation that is separate from background environmental microbiota present in nest material.

### Social interaction strength correlates with microbial sharing

Having established the relative contributions of colony differences and species-specific factors on microbiome structure, we asked to what extent social and trophic interactions with ants leave signatures in the core myrmecophile species’ microbiota. *A priori*, we hypothesized that *Sceptobius* and possibly *Myrmecophilus* may possess microbial communities most similar to that of the host ant due to their close physical interactions with workers. To test this hypothesis, we first filtered out sequences from putative intracellular endosymbionts in order to highlight the fraction of the microbiota most likely to be horizontally transferred during insect behavioral interactions. We also removed *Liometoxenus* from this analysis because only one sample from this rare species still met the minimum threshold of 1000 reads after removing the putative endosymbiont sequences. We then compared the distributions of pairwise distances and dissimilarity metrics calculated for each myrmecophile species with respect to the ant host. Samples were compared within nest sites, due to the aforementioned effect of colony identity on microbial community composition (**Fig. 2F**; **Table S2**).

By both Jaccard distance and Bray-Curtis dissimilarity, *Sceptobius* displayed the smallest median dissimilarity to *Liometopum*, followed in increasing order by *Myrmecophilus* and *Platyusa* (**Fig. 3A**, **Fig. S3**); for this analysis, all sample types except wash samples were considered together for each species (for example, *Liometopum* samples included whole body, dissected body, and gut samples). Thus, *Sceptobius* tended to possess the most similar non-endosymbiotic microbiota to that of its host ant, and *Platyusa* the least similar, regardless of whether bacterial relative abundances were taken into account. Notably, the same pattern was also observed for the body wash samples of surface microbiota: following removal of all ‘background’ environmental ASVs that were present in the nest material samples, we uncovered the greatest similarity in the surface microbiotas of *Sceptobius* and *Liometopum* (**Fig. 3B**). This is consistent with direct sharing of stably-associated external microbial symbionts between ants and socially integrated myrmecophiles—in this case we think driven by ant grooming, and phoretic attachment of the beetles to workers. The less pronounced but still physically close interactions between *Myrmecophilus* crickets and *Liometopum* workers also appear to provide routes for some amount of microbial transfer. Given that host ants outnumber myrmecophiles by two to three orders of magnitude per nest, it is unlikely that the specific worker ants we randomly sampled from colonies had recently—if ever—interacted with *Sceptobius* or *Myrmecophilus*. In contrast, every *Sceptobius* or *Myrmecophilus* individual within a nest is engaged in constant social interactions with ants. Hence, we propose that the shared microbes in the ant and myrmecophile samples derive from the ant, rather than the other way around.

To gain further resolution into this putative ant-myrmecophile pan-microbiome, we identified specific ASVs that were shared between specific sets of insect species. Kruskal-Wallis tests with discrete false discovery rate control (DS-FDR) (Jiang et al., 2017) revealed that 37 ASVs were differentially abundant across *Liometopum*, *Sceptobius*, *Myrmecophilus*, and *Platyusa* samples (**Table S3**). Among these 37 ASVs, 24 were shared by *Liometopum* and *Sceptobius*, while 21 were shared by *Liometopum* and *Myrmecophilus*. In contrast, only 10 ASVs were shared by *Liometopum* and *Platyusa* (**Fig. 3C**). For most of the ASVs shared between *Liometopum* and either *Sceptobius* or *Myrmecophilus*, the ASVs were more prevalent among the *Liometopum* samples, implying again that the directionality of microbial sharing tends to be from ant to myrmecophile rather than vice versa.

The results obtained for specific ASVs provide further evidence for microbial sharing between host ants and myrmecophiles that is correlated with the degree of social interaction between the insects. We subsequently determined how patterns of microbial sharing between *Liometopum* and the staphylinids differed by body part, by exploiting *Sceptobius* and *Platyusa*— the two species that respectively represent the most tightly and weakly integrated myrmecophiles in this study. Because *Sceptobius* physically interacts with ants using its mouthparts and legs (Danoff-Burg, 1996), engages in trophallaxis, and because staphylinids in general are capable of pre-oral digestion (Thayer, 2005), we hypothesized that the microbiota of *Sceptobius* heads and bodies may exhibit the greatest similarity to the microbiota of the ant. Consistent with this idea, pairwise Jaccard distances revealed that the microbiota of *Sceptobius* heads was indeed the closest among the myrmecophile body parts to ant bodies, followed in increasing order by *Sceptobius* bodies and guts (**Fig. 3D**). In contrast, *Platyusa* bodies, guts, and heads were all significantly less similar to the ant body microbiota than was any single body region of *Sceptobius* (**Fig. 3D**). These trends were also recovered in the Jaccard-distance-based NMDS ordination of ant, *Sceptobius*, and *Platyusa* samples marked by body part (**Fig. 3E**), and evident once more via Bray-Curtis dissimilarity (**Fig. S4A**). *Sceptobius* body parts were also significantly more similar than *Platyusa* body parts to ant gut samples (**Fig. S4B, C**).

That the microbiotas of *Platyusa* and *Liometopum* are more clearly distinct implies that their transient interactions, via chemical appeasement and predation, do not usually result in extensive microbial sharing. Interestingly however, using Bray-Curtis dissimilarity, *Platyusa* guts had a long tail of samples with relatively low dissimilarity to ant bodies and ant wash samples (**Fig. S4A, D**). This could possibly reflect consumption of host ant workers, but if so, we speculate that this similarity may reflect transient residents of the *Platyusa* gut rather than true symbionts. *Sceptobius* heads and bodies also displayed a tail of relatively low Bray-Curtis dissimilarities to ant wash samples (**Fig. S4D**). This observation is consistent with microbial sharing between *Liometopum* and *Sceptobius* driven at least in part by *Sceptobius* performing strigilation or latching onto the ant with its mouthparts, as well as by phoresy.

In addition to examining sharing of specific microbial ASVs, we employed weighted UniFrac to account for phylogenetic relationships between microbial strains present in the different insect species. Strikingly, weighted UniFrac distances suggested that *Sceptobius* body parts were even more similar to ant bodies and guts than was evident from Jaccard distances or Bray-Curtis dissimilarity. Indeed, by this metric, *Sceptobius* body parts were roughly as similar to ant bodies and ant guts as ant bodies and guts were to each other, whereas *Platyusa* body parts were still significantly more dissimilar to both ant bodies and guts (**Fig. S4E, F**). Given that UniFrac distances—unlike Jaccard distances or Bray-Curtis dissimilarity—incorporate phylogenetic information, this finding could potentially suggest an element of convergent evolution between the microbiotas of *Sceptobius* and *Liometopum*, on top of direct exchange of specific microbial species.

### Myrmecophile microbiota composition is not contingent on deeper evolutionary history

The independent evolution of myrmecophily in the three aleocharine rove beetles enabled us to ask how these species’ microbiotas were contingent on their evolutionary histories as opposed to their ecologies. The three beetles are inferred to have shared a ∼110 million year (MY)-old free-living common ancestor (Maruyama & Parker, 2017). Subsequently, *Sceptobius* likely diverged from its free-living sister, *Lissagria*, ∼50 MY ago, but the timing of its ensuing transition to life in *Liometopum* colonies is uncertain. For *Liometoxenus*, crown-group members of its tribe, Oxypodini, are inferred to have arisen in the Middle Eocene, implying that *Liometoxenus* is younger still and that its transition to myrmecophily probably occurred within the past 40 MY (Maruyama & Parker, 2017). Finally, the three lomechusines, *Drusilla*, *Pella*, and *Platyusa*, share a common ancestor ∼65 MY (Maruyama & Parker, 2017), implying a younger timeframe for *Platyusa’s* association with *Liometopum*. For this analysis, we initially included only the ASVs classified as putative intracellular endosymbionts, since these are primarily vertically transmitted (Bright & Bulgheresi, 2010; McCutcheon & Moran, 2012). Furthermore, sequences from such bacterial symbionts constituted the majority of reads for many of the beetle samples (**Fig. 2A, D**). If evolutionary relationships among the beetles were the main driving factor, due to ancient acquisition and vertical transmission of endosymbionts, then the myrmecophiles’ microbiotas may be more similar to closely related non-myrmecophilous sister groups than to each other. Conversely, if a myrmecophilous association with the same *Liometopum* host were a driving factor, then the myrmecophiles’ microbiotas may be more similar to each other.

To assess microbiota similarity across the different beetle species, we employed weighted UniFrac distances (Lozupone, Hamady, Kelley, & Knight, 2007), because we were interested in the phylogenetic relationships among the endosymbionts. We found that the two species with the most similar microbiotas by this measure were the *Liometopum* myrmecophiles *Sceptobius* and *Platyusa* (**Fig. 4A**). These species showed greater similarity to each other than to their outgroups, *Lissagria* and *Pella*+*Drusilla* respectively. However, beyond this relationship, an obvious pattern was not apparent: for example, the second closest pairing was composed of *Lissagria* and *Pella*, two phylogenetically distant outgroups, the former a free-living species and the latter a myrmecophile of *Lasius* ants (**Fig. 4A**). Furthermore, the microbiota of *Liometoxenus* (tribe Oxypodini) was closest to *Lissagria* (tribe Falagriini) rather than to either *Sceptobius* or *Platyusa* with which it coexists in the same host colony (**Fig. 4A**). Examination of the microbiotas of each staphylinid suggested that these patterns were largely driven by the small numbers of putative endosymbionts associated with each species (**Fig. 4B, C**). For example, both *Sceptobius* and *Platyusa* possessed only *Rickettsia*, while *Pella* and *Lissagria* possessed endosymbiont8 and members of the *Rickettsiaceae* family (*Rickettsia* in *Pella* and *Wolbachia* in *Lissagria*). *Liometoxenus* possessed only endosymbiont8, placing it closer to *Pella* and *Lissagria* than to *Sceptobius* or *Platyusa*. Finally, *Drusilla* only possessed *Rickettsiella* (a member of the Gammaproteobacteria and hence more closely related to endosymbiont8 than to *Rickettsia* or *Wolbachia*).

Evidently, the endosymbiotic microbial communities of the three aleocharine members of the core myrmecophile community cannot be explained by the evolutionary histories of these beetle species. This qualitative lack of phylosymbiosis—co-cladogenesis between microbial communities and host organisms—suggests that these putative intracellular endosymbionts were either acquired after the branching of the different myrmecophiles from outgroups and from each other, and/or were lost over time in particular lineages. There was likewise no clear effect of lifestyle on the propensity to possess particular endosymbiont taxa. For example, while *Sceptobius* and *Platyusa* shared relatively closely related endosymbionts (different *Rickettsia* ASVs), *Liometoxenus*, which lives with the same host ant species, possessed a very different set of species. And while *Pella* and *Lissagria* possessed phylogenetically similar endosymbionts, one is a myrmecophile while the other is free-living. Thus, although our data do not rule out the possibility that some of the detected putative endosymbionts are a consequence of (or are adaptive for) a myrmecophilous lifestyle, the acquisition and/or loss of particular clades of endosymbionts in these staphylinid species may primarily represent chance evolutionary events within each separate beetle lineage.

Finally, we assessed whether there was any evidence for a primary role of ecology versus evolutionary history among the insects with regard to shaping the composition of their non-endosymbiont microbiotas. For this analysis, after removing putative intracellular endosymbiont sequences from the data, we focused on six of the insect species: *Liometopum, Sceptobius, Lissagria, Platyusa, Pella*, and *Lasius.* We hypothesized that if ecology were the primary driver of the non-endosymbiont microbiotas, the microbiotas of *Platyusa* and *Pella* would be more similar to that of their ant hosts (*Liometopum* and *Lasius*, respectively) than to that of each other. Likewise, the microbiota of *Sceptobius* may be more similar to that of *Liometopum* than to that of its free-living relative, *Lissagria*. Conversely, if the impact of evolutionary history were the primary driver, *Pella* and *Platyusa* would be more similar to each other than to their ant hosts, and *Sceptobius* might be more similar to *Lissagria* than to *Liometopum*. Intriguingly, weighted UniFrac distances revealed a trend that was a hybrid of these two scenarios (**Fig. S5**). *Pella* and *Platyusa* were indeed more similar to each other than either were to their respective ant hosts, and *Lissagria* and *Sceptobius* were also more similar to each other than *Lissagria* was to *Liometopum* (which is found in the same region and environments as *Lissagria*, even though the two are not behaviorally associated), albeit these trends were not statistically significant. These observations suggest a possible influence of shared evolutionary history on the non-endosymbiont microbiota of these beetles. Nevertheless, the closest of all the pairings was *Sceptobius* and *Liometopum*, suggesting that strong ecological and behavioral ties between highly divergent insect species can override this potential phylogenetic effect.

### Genomic insights into the *Sceptobius* intracellular endosymbiont

Adaptive functions of intracellular endosymbionts are well known in many insect species with specialized ecologies (Douglas, 1998; McCutcheon, 2021; Zientz et al., 2004). Whether such endosymbionts might play comparable roles in enabling obligate socially parasitic lifestyles, in which the insect has sacrificed any capacity for a free-living existence, is, however, unknown. Such lifestyles commonly involve extreme specialization for life inside host colonies, manifesting in multiple dimensions of the phenotype including behavior, chemical communication, reproductive physiology, diet and feeding strategy, as well as morphology. *Sceptobius* represents a virtuoso example of this type of organism, providing an opportunity to explore potential microbial involvement in its obligate dependence on *Liometopum* ants. We were struck by the high abundance of a single *Rickettsia* ASV in all *Sceptobius* samples, at frequencies of up to 95% of 16S reads per beetle. This bacterium occurred in all beetles of both sexes (importantly, *Sceptobius* shows no noticeable sex ratio distortion in the populations sampled for this study). The extreme dominance of this *Rickettsia* species was paired with an absence of other possible contenders for primary endosymbionts.

To shed light on this bacterium—herein named “RiSlat” for “*Rickettsia* endosymbiont of *Sceptobius lativentris*”—we assembled its genome using Illumina short reads obtained from genomic DNA extracted from a single male *Sceptobius* body, combined with Oxford Nanopore long reads from pooled beetles (see Materials and Methods). The final genome assembly consisted of two contigs with a total size of 1,597,619 bp, and with an average GC content of 32%, typical of Rickettsial genomes (**Table S4**). The completion score was over 99% (**Fig. 5B**) suggesting that our assembly is a near complete genome. Genomic annotation identified 1825 protein coding sequences with an average size of 741 bp accounting for coding density of circa 85%. Out of the 1825 predicted protein-coding genes, 510 (∼30%) were annotated as hypothetical without known functional domains, while 294 genes (∼16%) were predicted to code for transposases (**Table S4**).

**Figure 5.**
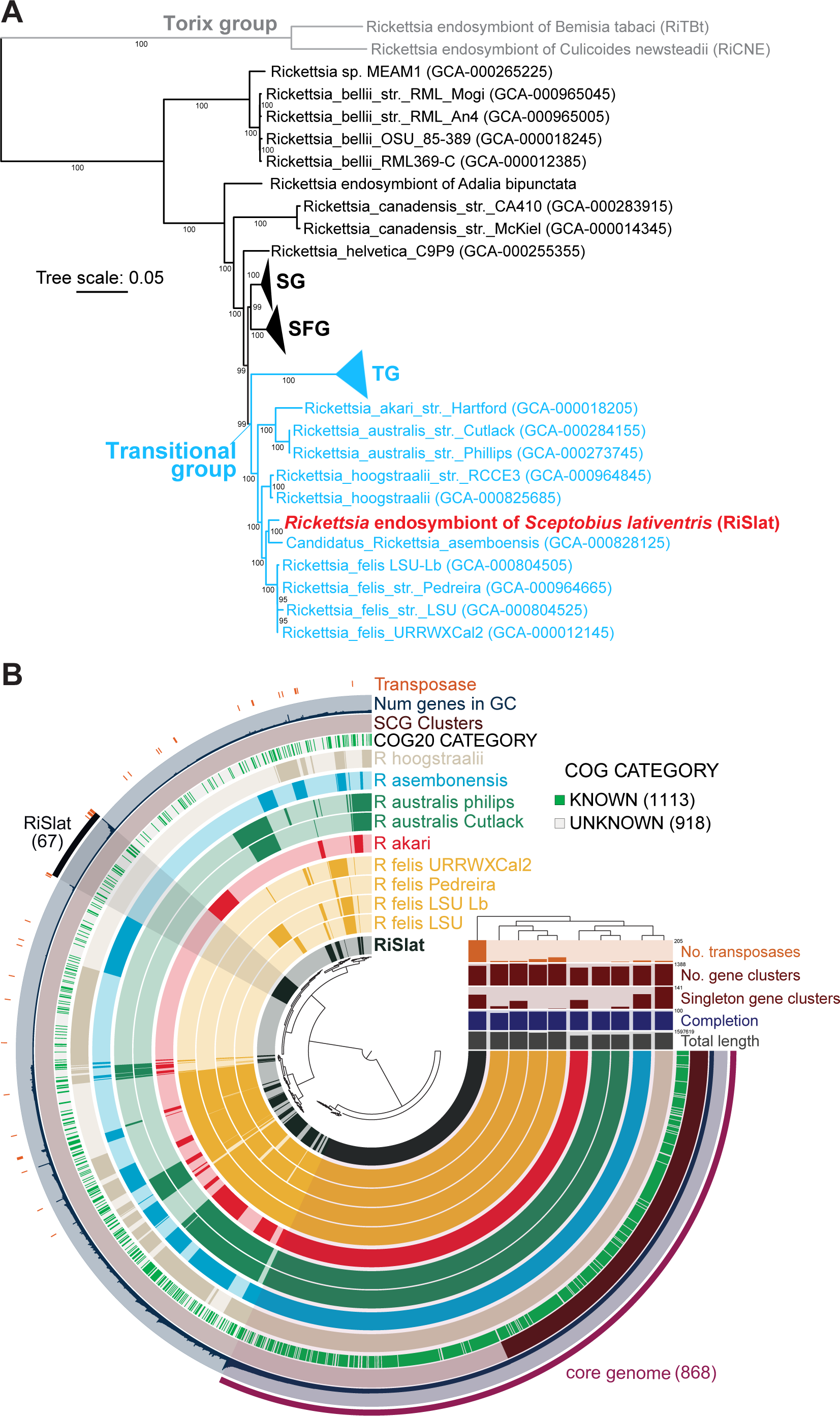
Comparative genomics of the *Rickettsia* endosymbiont of *Sceptobius lativentris*. **A:** Phylogenomic position of *Rickettsia* endosymbion**t** of *Sceptobius lativentris* (RiSlat) based on concatenated analysis of 267 single copy core proteins. Relationships were inferred using the maximum likelihood criterion as implemented in IQTREE 2.0.3 (Minh et al., 2020) under the JTT+F+R5 model. Numbers on branches represent support values based on 1000 ultrafast bootstrap replicates. Some of the *Rickettsia* groups have been collapsed for visualization purposes. SG: Scapularis group; SFG: Spotted fever group; TG: Typhus group. **B:** Comparative genomic analysis across members of the Transitional group of Rickettsia using anvi’o package (Eren et al., 2015). Each of the ten inner layers represent the gene clusters (GC) identified in the ten Rickettsia genomes and ordered based on their distribution across the ten genomes. Rickettsia genomes were clustered and color coded based on gene cluster presence absence (dark color and light color represent presence and absence of a gene cluster). The core genome, the single core genome (SCG) and the gene clusters present only in the *Rickettsia* endosymbiont of *Sceptobius lativentris* (RiSlat) are indicated. Number in the parenthesis correspond to number of gene clusters.

The phylogenetic position of the RiSlat in relation to known *Rickettsia* groups was established based on a concatenated set of 267 single core proteins shared between 86 publicly available *Rickettsia* genomes. Maximum likelihood analysis placed RiSlat within the “transitional group” of *Rickettsia*, most closely associated to *Ca*. Rickettsia asemboensis (**Fig. 5A**). The transitional group is known to host both pathogenic strains, such as *Rickettsia felis*, the causative agent of a typhus-like flea-borne rickettsiosis (Azad, Radulovic, Higgins, Noden, & Troyer, 1997), as well as obligate mutualistic endosymbionts including the *Rickettsia* symbiont of the parthenogenetic booklouse, *Liposcelis bostrychophila* (Hagimori, Abe, Date, & Miura, 2006; Perotti, Clarke, Turner, & Braig, 2006; Yusuf & Turner, 2004). We performed a comparative genomic analysis across members of the transitional group to identify shared and unique features potentially associated with host specialization and biological function. A total of 2410 gene clusters were identified among the ten available genomes belonging to the transitional group. 868 of the gene clusters were shared between all genomes, representing the core genome of the transitional group. Overall, the RiSlat genome shared many of the typical features associated with the *Rickettsia* genus including a type-IV secretion system, 11 genes coding for putative surface antigens, and several genes coding for components of toxin-antitoxin systems (14 toxins and 17 antitoxins). We identified 67 gene clusters comprising a total of 476 genes specifically associated with the RiSlat genome (**Fig. 5B**). 166 of these genes had at least one homolog in another *Rickettsia* genome outside of the transitional group, while 310 genes appear to be uniquely associated with RiSlat (**Table S4**).

(Minh et al., 2020)(Eren et al., 2015)Among the RiSlat-specific loci were 113 (∼36%) genes with no assigned function, a toxin-antitoxin pair, and loci encoding a leucine-rich repeat-containing protein and a D-Ala-D-Ala carboxypeptidase. Most notably, however, 193 genes (∼62%) were annotated as putative transposases belonging to the IS1380 family with a putative gamma-proteobacterial origin. This large expansion of transposable elements is markedly higher than in other members of the transitional group (**Fig. 5B**). The majority of these transposases share more than 95% similarity, suggesting a relatively recent or even an ongoing expansion. Despite their high number and similarity, we did not detect significant synteny breaks between RiSlat and other genomes from the transitional group as a result of homologous recombination (**Fig. S6A-D**). Furthermore, the fraction of interrupted genes in RiSlat was estimated to be only ∼13.7% (128 genes with length <80% of their top hit in Swiss-Prot database), which is lower than that of other members of the transitional group (**Fig. S6E**). This suggests non-significant pseudogenization despite extensive proliferation of transposable elements.

Distant homologs of loci associated with cytoplasmic incompatibility in *Wolbachia* have been reported in some *Rickettsia* (including the transitional group (Gillespie et al., 2018), but we find that these are not present in the RiSlat genome. Finally, we assessed the metabolic potential of the RiSlat genome and compared it to other members of the transitional group by estimating the completeness of metabolic pathways including carbohydrate and energy metabolism, lipid and glycan metabolism, amino acid metabolism, and metabolism of cofactors and vitamins (**Fig. S7**). The metabolic potential of RiSlat resembles that of other members of the transitional group.

Both cofactors and B vitamin biosynthesis are limited, while most of the amino acid biosynthesis pathways are absent or incomplete. These results tentatively argue against a role for RiSlat as a nutritional symbiont, and instead indicate that this bacterium may play an alternative role in the biology of its host beetle.

## Discussion

We have characterized microbiotas associated with a host ant species and its community of symbiotic, socially integrated myrmecophiles. Our results shed light on how obligate behavioral relationships between different animal species can affect their associated microbial communities. We found that the microbiotas of the host ant and its four core myrmecophiles each display unique, species-specific characteristics, ranging from the level of taxonomic diversity to the presence of putative intracellular endosymbionts. Nevertheless, the similarity of the microbiotas of the different myrmecophiles to that of their common host ant was qualitatively correlated with each species’ level of social interaction with the latter. Moreover, we detected multiple specific bacterial species that were shared between highly interacting species (e.g. *Liometopum* and *Sceptobius*) but not between more weakly interacting species (e.g. *Liometopum* and *Platyusa*). We further identified body part-specific patterns of microbial sharing that match known aspects of the behavioral interactions between these myrmecophiles species and their host ant. The patterns of sharing we have uncovered provide evidence for a cross-species pan-microbiome emerging from close physical relationships between these insect species. Such behaviors include phoretic attachment to workers, grooming, trophallaxis, and strigilation. Analogues of these behaviors have evolved convergently in many other ant-myrmecophile relationships, some of which rank among the most intimate and specialized forms of animal-animal symbiosis (Hölldobler & Wilson, 1990; Kistner, 1979, 1982; Maruyama et al., 2009; Parker, 2016). We predict that analogous pan-microbiotas may emerge generally in these contexts of exceptionally close physical associations between socially integrated myrmecophiles and their hosts.

Our findings add to a growing body of evidence that microbial exchange is a pervasive feature of socially complex metazoan interspecies interactions (Pringle & Moreau, 2017; Song et al., 2013). To our knowledge, our study is the first to reveal a correlation between the strength of the social interaction with a given partner (in this case, *Liometopum*), and the degree of microbial sharing. Whether microbial sharing serves an adaptive function in the context of myrmecophily is presently unclear. Previous studies have shown how experimentally altering the surface microbiome of red harvester ants (*Pogonomyrmex barbatus*) leads to increased aggression from nestmates (Dosmann, Bahet, & Gordon, 2016), implying that ants can either detect differences in microbial communities, or that microbes modulate the pheromonal chemistry of workers, perhaps by altering their CHC profiles. Microbial exchange may thus be critical for myrmecophiles such as *Sceptobius*, which rely on chemical resemblance for social integration, with grooming and close physical contact enabling horizontal acquisition of both CHCs and surface microbes. The presence of host ant-derived bacteria in the head of *Sceptobius* may aid in pre-oral digestion (Thayer, 2005), or be a consequence of strigilation over the ant body surface. Alternatively, it may arise from the beetle grasping the ant via its mandibles—a stereotyped behavior that appears to anchor *Sceptobius* to the worker, freeing up its legs for grooming (**Video S1**).

Our findings contrast with a recent survey of microbial diversity in beetles associated with the red wood ant, *Formica polyctena,* in Europe (Kaczmarczyk-Ziemba, Zagaja, Wagner, Pietrykowska-Tudruj, & Staniec, 2020). Here, no evidence of microbial sharing was found, but all the beetle species examined in the study are unspecialized, non-socially integrated taxa that exhibit no physical interactions with their host ant, perhaps further emphasizing the importance of behavioral intimacy in establishing a cross-species microbiota. As in our study, Kaczmarczyk-Ziemba et al (2020) found a high prevalence of putative intracellular endosymbionts among the staphylinid species. By incorporating multiple rove beetle species that span a range of phylogenetic relationships and interactions with ants, including outgroup species, we were able to investigate whether either lifestyle or phylogeny exert a strong influence on these intracellular bacteria. We conclude that there is little phylogenetic signal in this component of the staphylinid microbiotas, with little evidence for phylosymbiosis. Although two closely related species, *Platyusa* and *Pella*, shared the same *Rickettsia* variant (ASV7), other instances of shared endosymbiont ASVs were between much more distantly related species, such as *Pella* and *Sceptobius* (ASV1), or *Pella* and *Lissagria* (ASV3). We also found no clear correlation between overall intracellular microbiota composition and degree of myrmecophily or the identity of the host ant species. Together, these results suggest that intracellular endosymbionts, which appear to dominant the rove beetle microbiota, may have largely been acquired through chance events in their evolutionary history, rather than reflecting clade-specific physiological factors or selective pressures imposed by myrmecophily. The lack of conserved taxonomic characteristics among the different myrmecophile microbiotas does not, however, preclude functional convergence. In the future, shotgun metagenomic sequencing might reveal whether the latter is indeed the case among myrmecophiles.

Insect endosymbionts are often vertically transmitted, but the lack of a strong phylogenetic signal in the microbiotas of these staphylinids is perhaps not surprising given that members of *Rickettsia* and *Wolbachia* (and presumably also *Rickettsiella* and the “endosymbiont8” clade) are typically facultative endosymbionts. Such bacterial taxa are often not required for the survival of their hosts, and are consequently prone to multiple interspecies horizontal transfers and losses over evolutionary time (Moran et al., 2008; Perlman, Hunter, & Zchori-Fein, 2006). The apparent lack of long-term microbial associations in rove beetle contrasts with other insects where endosymbionts perform vital metabolic functions for their host insects, the two organisms consequently engaging in pronounced phylosymbiosis (Ivens, Gadau, Kiers, & Kronauer, 2018). Despite the comparative recency of their associations with rove beetles, the high within-species prevalence of the most abundant intracellular bacterial species in our study suggests that some have likely become established as stable, vertically transmitted endosymbionts. The exact anatomical locations of these bacterial species remain to be determined, but their broad distributions can be inferred from high abundances across different dissected beetle body parts. Possible physiological impacts of these microbes are presently unknown, but facultative endosymbionts generally fall into two categories: those that directly manipulate reproductive outcomes for their host (e.g. through male killing or cytoplasmic incompatibility with uninfected mates), and those that offer a mutualistic benefit to their host and therefore indirectly increase the host’s reproductive output (Moran et al., 2008).

The high bacterial load and ubiquity of RiSlat in members of both sexes of the *Sceptobius* population hints at a potentially beneficial role in the biology of its myrmecophile host. RiSlat possesses unique genomic features, in particular a large expansion of IS elements of g-proteobacterial origin, which are not found in other *Rickettsia*. Metabolically, however, RiSlat largely resembles other members of the transitional group of *Rickettisa*. Importantly, we find no evidence of nutritional provisioning to the beetle—a phenomenon that has arisen recurrently across the Coleoptera, in species that feed on nutrient-poor or recalcitrant diets. Microbial supplementation of tyrosine, which is necessary for formation of the heavily sclerotized beetle cuticle (Noh, Muthukrishnan, Kramer, & Arakane, 2016), appears in particular to be a relatively common service provided by beetle endosymbionts (e.g. Anbutsu et al., 2017; Engl et al., 2018; Vigneron et al., 2014). Although the diet of *Sceptobius* is not completely known, it likely includes trophallaxis with host ants and predation on ant brood (Danoff-Burg, 1996)—both of which represent protein-rich trophic resources. Hence, despite the highly specialized biology of *Sceptobius*, we posit that an obligate metabolic dependency on an endosymbiotic bacterium may not be of relevance, at least during the adult stage.

If RiSlat does benefit host biology, we speculate that an alternative role may be as a defensive symbiont—a function previously ascribed to some non-pathogenic *Rickettsia* endosymbionts of other insect hosts (Hendry, Hunter, & Baltrus, 2014; Łukasik, Guo, Asch, Ferrari, & Godfray, 2013). The precedent for endosymbionts playing a role in host defense strategies is particularly intriguing, given that myrmecophilous beetles coexist with aggressive and chemically defended ants, colonies of which are themselves targeted by a huge diversity of pathogens and parasites (Bekker, Will, Das, & Adams, 2018; Kistner, 1982; Wojcik, 1989). Although we detected no consistent correlation between possessing a particular endosymbiont and having a myrmecophilous lifestyle, different microbial taxa could potentially fulfill similar functions in different myrmecophiles. Future experiments, including antibiotic treatments to reduce or eliminate the endosymbiont load, may reveal how these endosymbionts might contribute to the beetles’ fitness in the context of the ant society.

## Acknowledgments

We thank Clive Turner (UK) for providing living specimens of *Pella cognata* and *Lasius fuliginosus*, Tim Struyve (Belgium) for providing *Drusilla canaliculata* and Julian Wagner (Caltech) for *Lissagria laeviuscula*. Preliminary stages of this work were facilitated by a seed grant from Caltech’s Center for Environmental Microbial Interactions, and we greatly appreciate the support and generosity of Dianne Newman (Caltech) throughout the course of this study. This work was funded by an Army Research Office MURI award, W911NF1910269, to J. Parker.

## Supplemental figure legends

**Figure S1.**
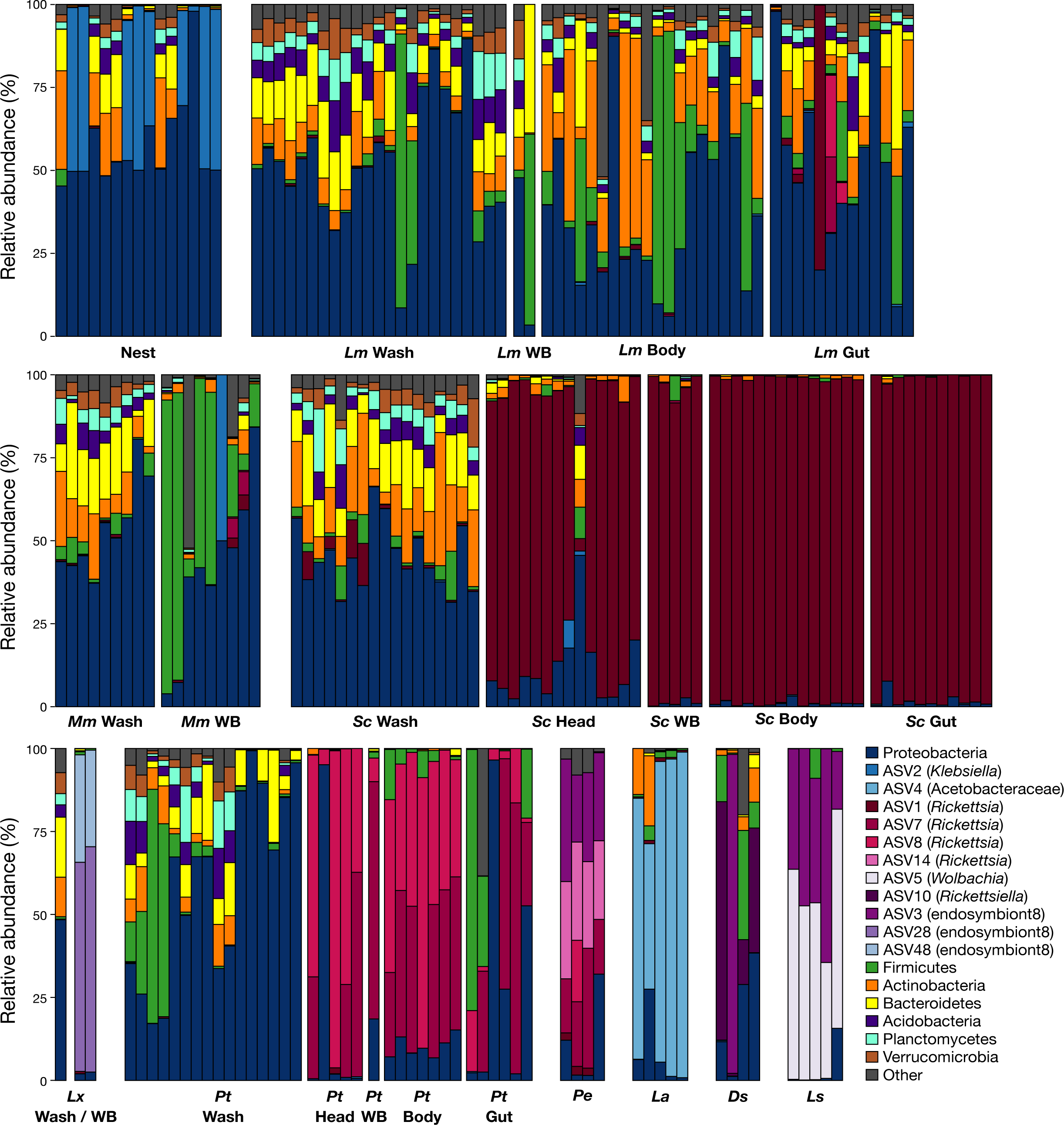
Bar plots showing relative abundance of different bacterial taxa in individual samples of the studied species. *Lm* = *Liometopum* (ant), *Mm* = *Myrmecophilus* (cricket), *Sc* = *Sceptobius* (beetle), *Lx* = *Liometoxenus* (beetle), *Pt* = *Platyusa* (beetle), *Pe* = *Pella* (beetle), *La* = *Lasius* (ant), *Ds* = *Drusilla* (beetle), *Ls* = *Lissagria* (beetle), WB = whole body.

**Figure S2.**
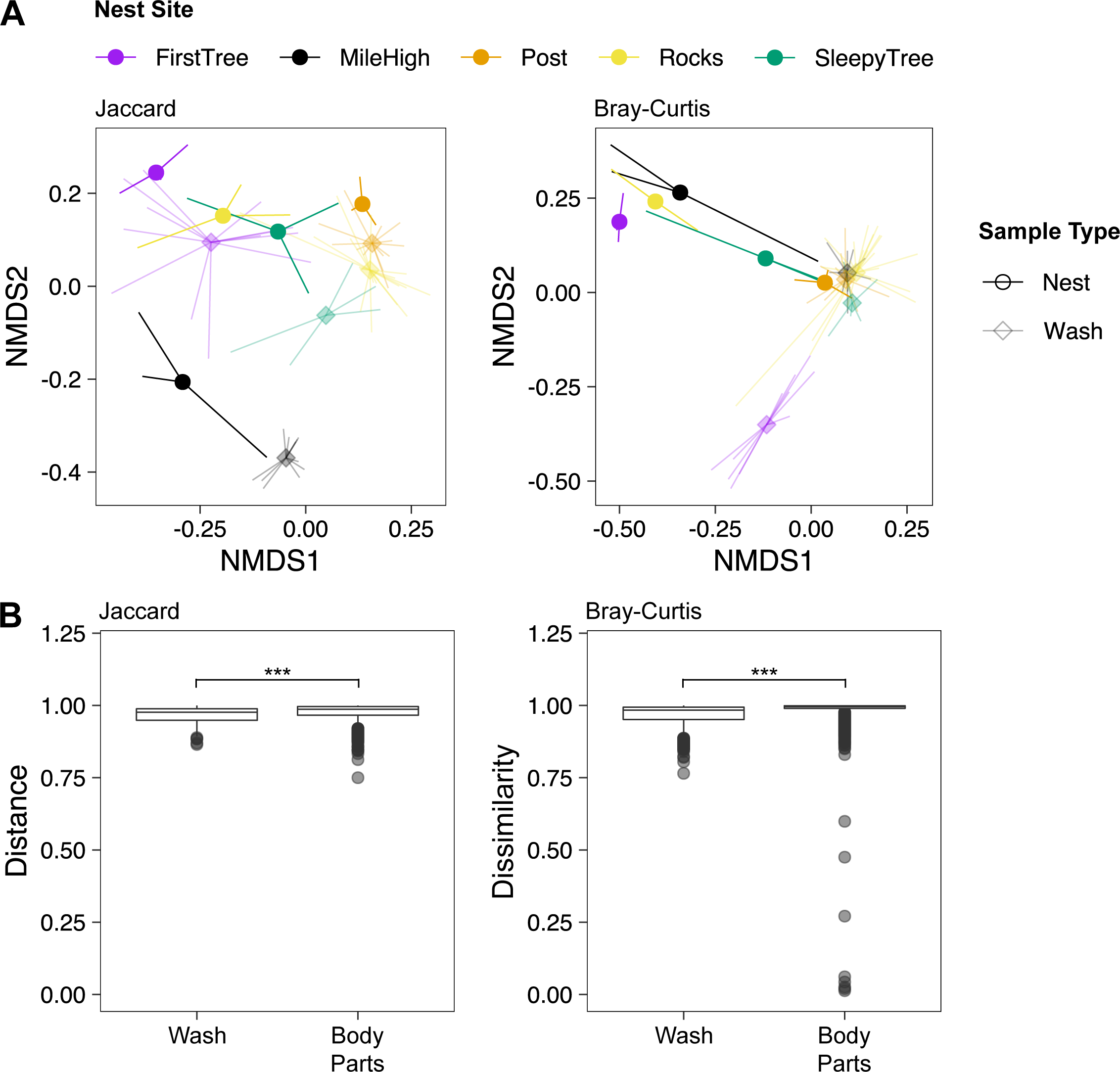
**A:** NMDS ordination of nest and wash samples, based on Jaccard distance or BrayCurtis dissimilarity and colored by nest site. Note that in some cases, individual samples are obscured by their proximity to each other or to the group centroid. **B:** Pairwise Jaccard distances and Bray-Curtis dissimilarities for wash samples and body part samples versus nest samples.

**Figure S3.**
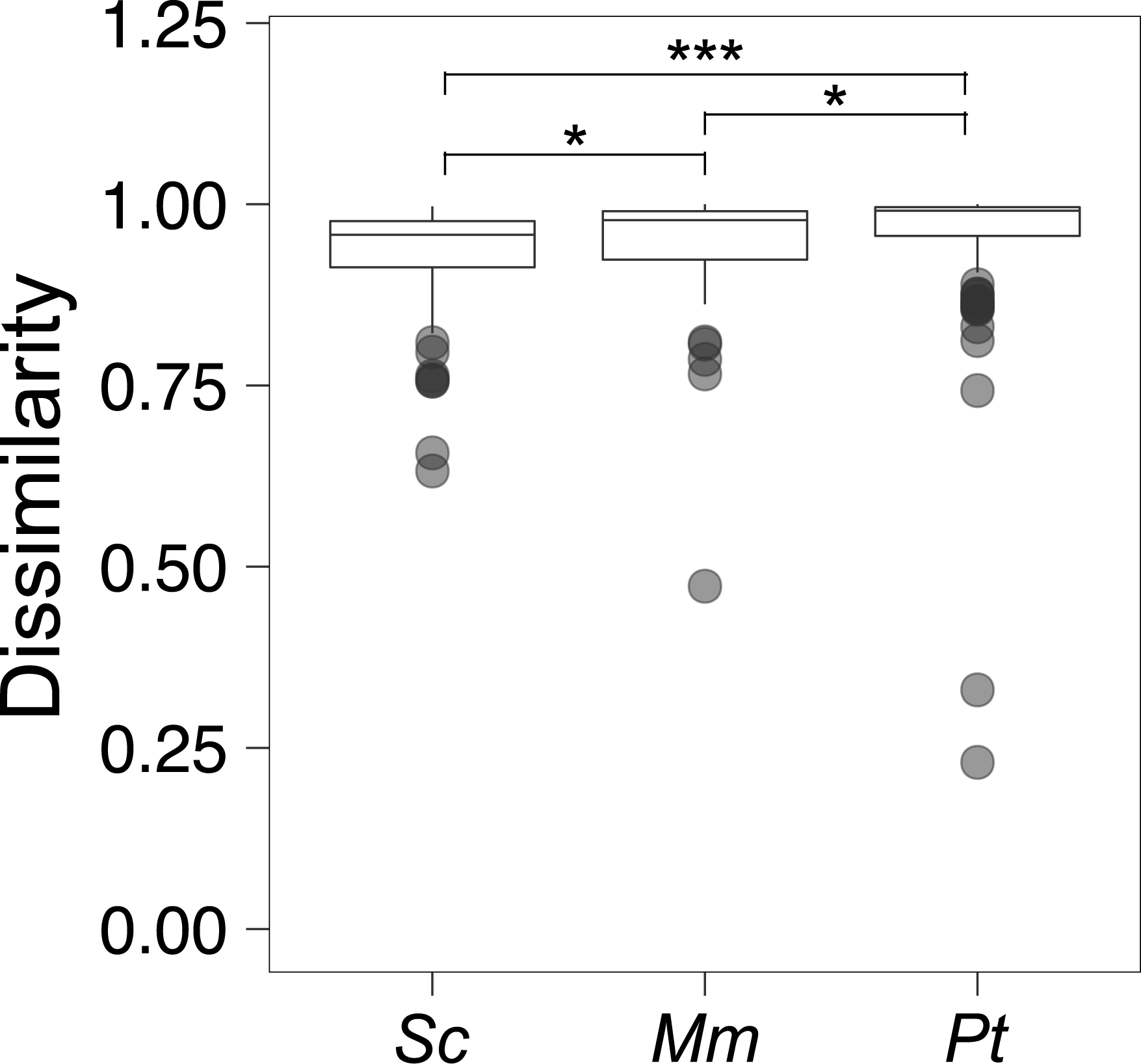
Bray-Curtis pairwise dissimilarities for *Sceptobius* (*Sc*), *Platyusa* (*Pt*), and *Myrmecophilus* (*Mm*) versus *Liometopum,* within nest sites, without endosymbionts. All sample types except wash samples (i.e. whole body samples in addition to dissected body parts where applicable) were included for each species.

**Figure S4.**
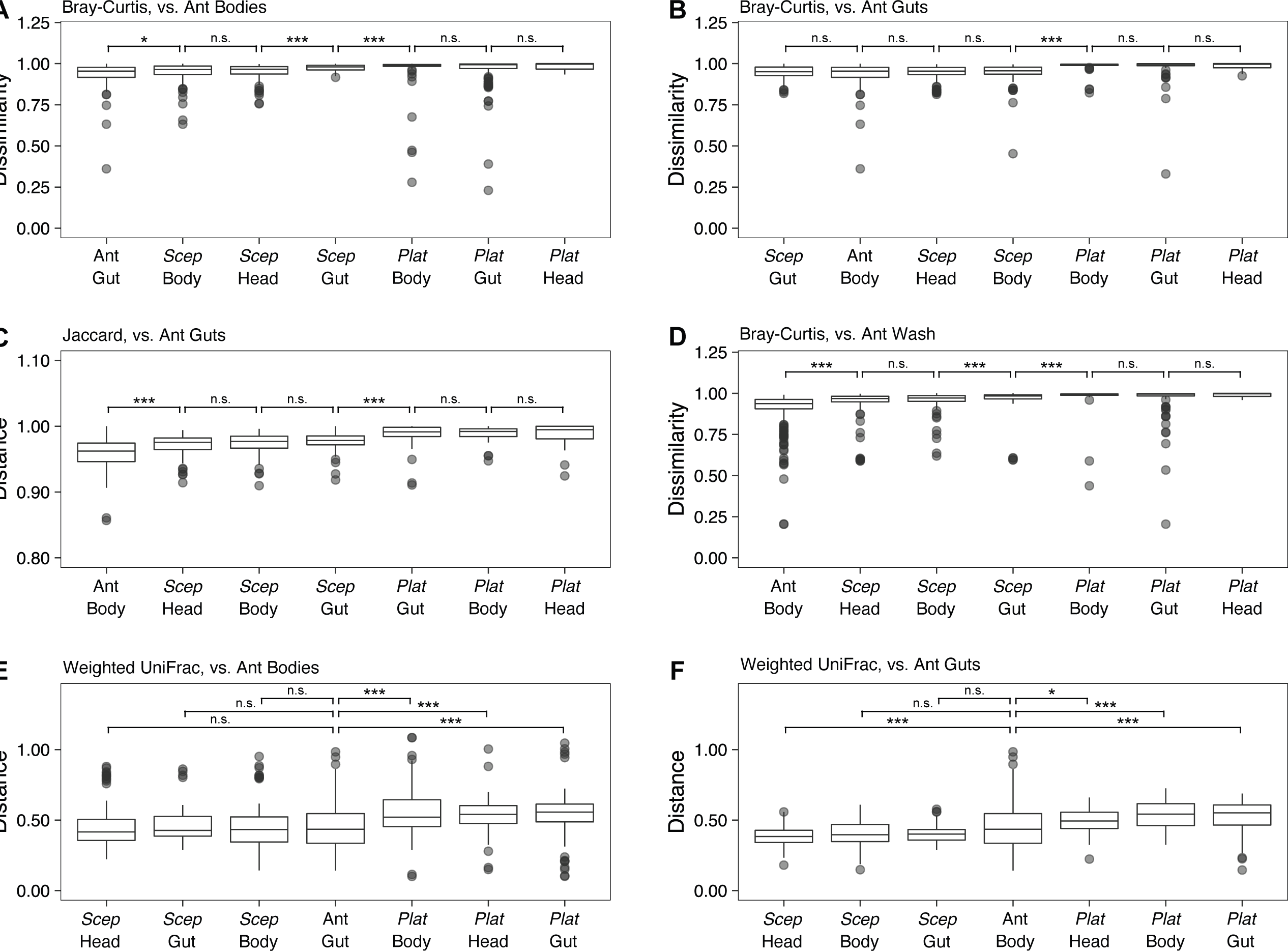
Pairwise distances or dissimilarities between different myrmecophile body parts and ant body parts, without endosymbionts. **A:** Bray-Curtis dissimilarities versus ant bodies. **B:** Bray-Curtis dissimilarities versus ant guts. **C:** Jaccard distances versus ant guts. **D:** Bray-Curtis dissimilarities versus ant wash samples. **E:** Weighted UniFrac distances versus ant bodies. **F:** Weighted UniFrac distances versus ant guts.

**Figure S5.**
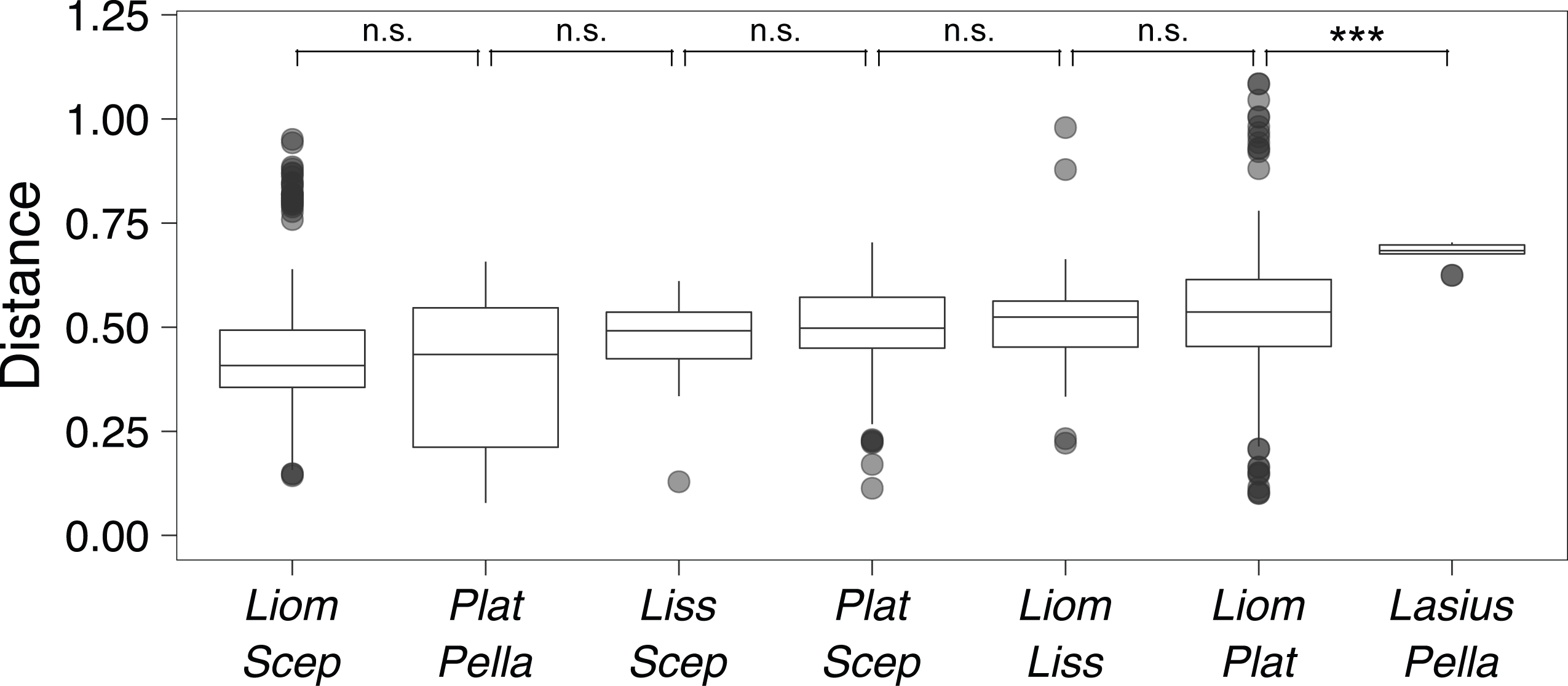
Pairwise weighted UniFrac distances between different staphylinids and ant species, without endosymbionts. All sample types were included except for wash samples.

**Figure S6.**
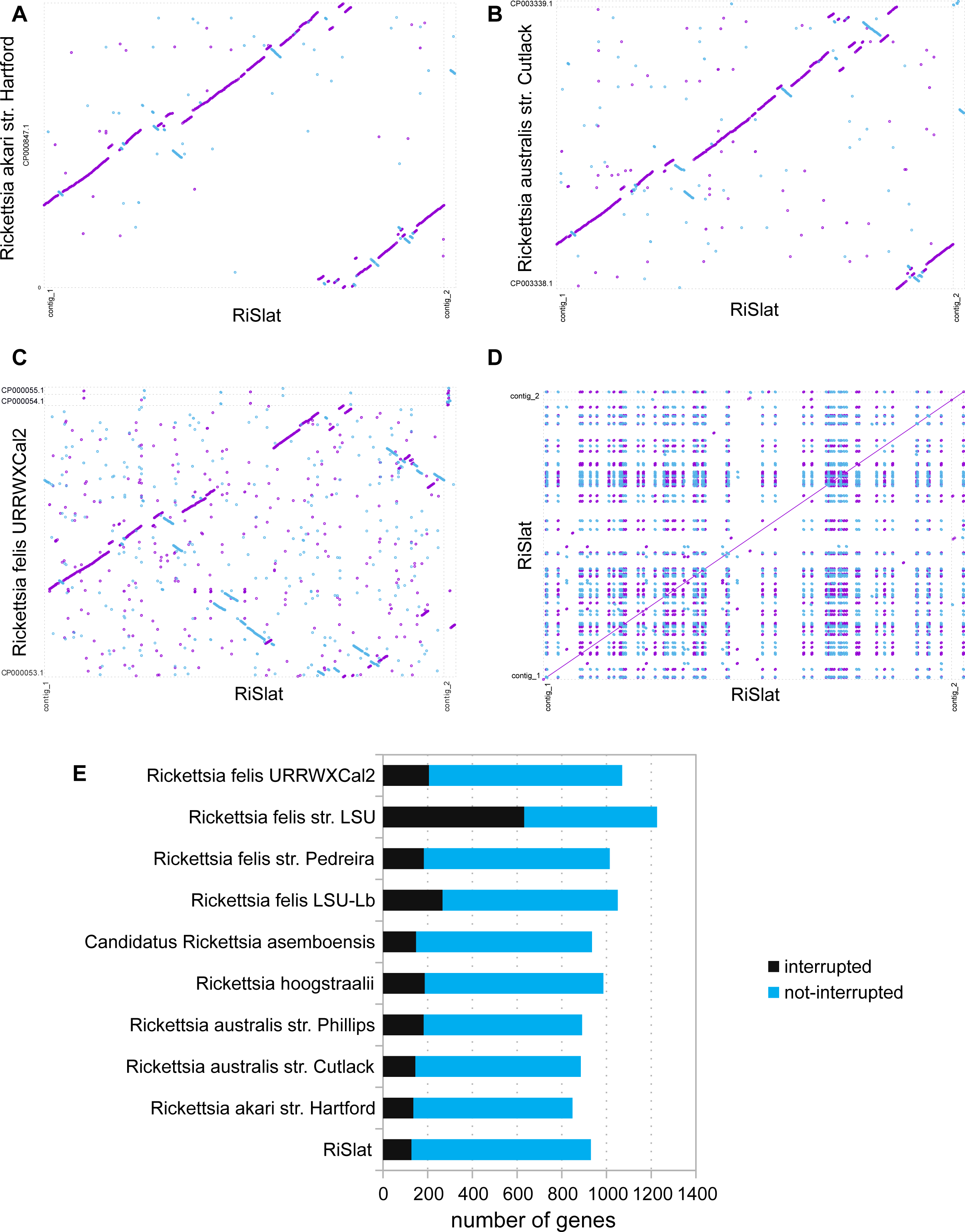
**A-C:** Syntenic comparison between RiSlat and related and complete genomes from the transitional group of *Rickettsia*. **D:** Mummer self-plots representing the repeat density (≥1500bp and ≥95% similarity) in the RiSlat genome. Magenta represents forward matches and blue represents reverse matches. **E:** Fraction of interrupted genes in RiSlat genome in comparison to other genomes from transitional group.

**Figure S7.**
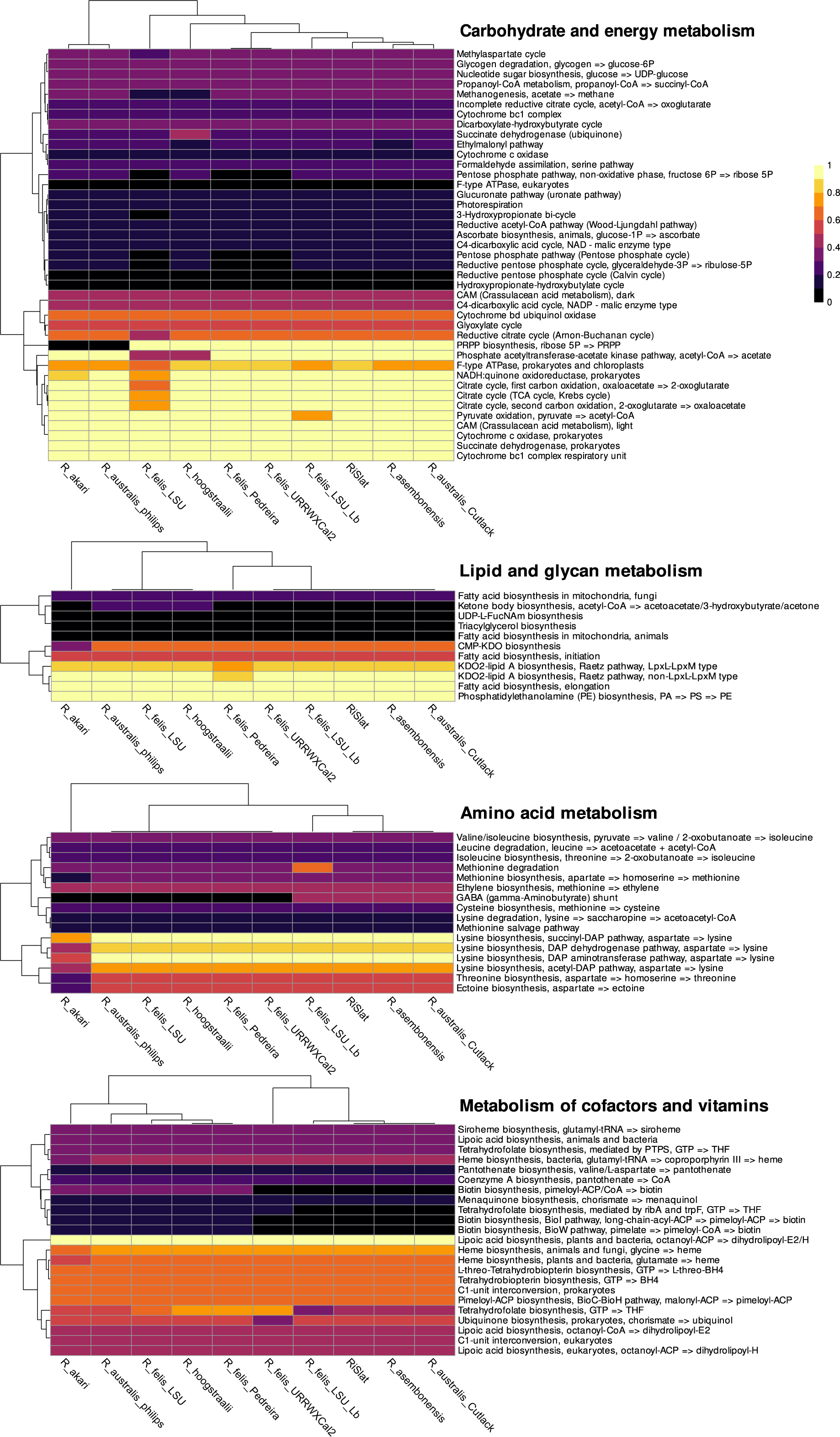
Reconstruction and comparative analysis of the metabolic potential across Transitional group Rickettsia. Pathway completeness for different KEGG functional categories were estimated using the “anvi-estimate-metabolism” program from the anvi’o package.

## Supplemental Videos

**Video S1. A** *Sceptobius lativentris* beetle mounted on top of *a Liometopum occidentale* worker. The beetle grasps the ant’s antenna in its mandibles and uses its tarsi to groom the ant’s body surface before smearing its tarsi over its own body. Grooming behavior transfers cuticular hydrocarbon pheromones from the ant onto the beetle. Video shot under infrared illumination at 90 frames per second.

## Supplemental Tables

**Table S1.** The frequency of each ASV per sample (normalized to 1000 reads). Mean frequencies per sample type, as well as sample metadata, are provided as separate tabs within this table.

**Table S2.** Summary of statistical tests used in this paper.

**Table S3.** Kruskal-Wallis test statistics and raw *p*-values for the 37 ASVs that were identified as differentially abundant across *Liometopum*, *Sceptobius*, *Myrmecophilus*, and *Platyusa* after using DS-FDR to control the false discovery rate. The order of the rows is the same as for the heatmap shown in **Fig. 3C**.

**Table S4.** Genome features of the *Rickettsia* endosymbiont of *Sceptobius lativentris* (RiSlat).

|  |  |
| --- | --- |
| Total size (bp) | 1,597,619 |
| Number of contigs | 2 |
| Largest contig (bp) | 1,551,323 |
| Average coverage per contig | contig_1: 85x, contig_2: 161x |
| GC contents (%) | 32 |
| Number of predicted CDS | 1825 |
| Average CDS length (bp) | 741 |
| Number of rRNAs | 3 |
| Number of tRNAs | 34 |
| Plasmids | Contig_2 is a possible low copy plasmid |

**Table S5.** Genome annotation file for the *Rickettsia* endosymbiont of *Sceptobius lativentris* (RiSlat).

## Supplemental Files

**File S1.** Fasta file of contigs 1 and 2 of the genome of the *Rickettsia* endosymbiont of *Sceptobius lativentris* (RiSlat).

**File S2.** GenBank flat file with complete genome annotation of the *Rickettsia* endosymbiont of *Sceptobius lativentris* (RiSlat).

## References

1. Akino, T. (2002). Chemical camouflage by myrmecophilous beetles Zyras comes (Coleoptera: Staphylinidae) and Diartiger fossulatus (Coleoptera: Pselaphidae) to be integrated into the nest of Lasius fuliginosus (Hymenoptera: Formicidae). Chemoecology, 12(2), 83–89.

2. Akre, R. D., & Hill, W. B. (1973). Behavior of Adranes taylori, a myrmecophilous beetle associated with Lasius sitkaensis in the Pacific Northwest (Coleoptera: Pselaphidae; Hymenoptera: Formicidae). Journal of the Kansas Entomological Society, 46, 526–536.

3. Anbutsu, H., Moriyama, M., Nikoh, N., Hosokawa, T., Futahashi, R., Tanahashi, M., … Fukatsu, T. (2017). Small genome symbiont underlies cuticle hardness in beetles. Proceedings of the National Academy of Sciences, 114(40), E8382–E8391. doi: 10.1073/pnas.1712857114

4. Azad, A. F., Radulovic, S., Higgins, J. A., Noden, B. H., & Troyer, J. M. (1997). Flea-borne rickettsioses: ecologic considerations. Emerging Infectious Diseases, 3(3), 319–327. doi: 10.3201/eid0303.970308

5. Bagnères, A.-G., Blomquist, G. J., Bagnères, A.-G., & Lorenzi, M. C. (2010). Chemical deception/mimicry using cuticular hydrocarbons. In Cambridge Core (pp. 282–324). Cambridge: Cambridge University Press. doi: 10.1017/cbo9780511711909.015

6. Beeren, C. von, Blüthgen, N., Hoenle, P. O., Pohl, S., Brückner, A., Tishechkin, A. K., … Kronauer, D. J. C. (2021). A remarkable legion of guests: Diversity and host specificity of army ant symbionts. Molecular Ecology. doi: 10.1111/mec.16101

7. Beeren, C. von, Brueckner, A., Maruyama, M., Burke, G., Wieschollek, J., & Kronauer, D. J. C. (2018). Chemical and behavioral integration of army ant-associated rove beetles - a comparison between specialists and generalists. Frontiers in Zoology, 15(1). doi: 10.1186/s12983-018-0249-x

8. Beeren, C. von, Schulz, S., Hashim, R., & Witte, V. (2011). Acquisition of chemical recognition cues facilitates integration into ant societies. BMC Ecology, 11(30), 1–12. doi: 10.1186/1472-6785-11-30

9. Bekker, C. de, Will, I., Das, B., & Adams, R. M. (2018). The ants (Hymenoptera: Formicidae) and their parasites: effects of parasitic manipulations and host responses on ant behavioral ecology. Myrmecological News, 28.

10. Birtel, J., Walser, J.-C., Pichon, S., Bürgmann, H., & Matthews, B. (2015). Estimating Bacterial Diversity for Ecological Studies: Methods, Metrics, and Assumptions. PLoS ONE, 10(4), e0125356. doi: 10.1371/journal.pone.0125356

11. Bologna, M. A., & Pinto, J. D. (2001). Phylogenetic studies of Meloidae (Coleoptera), with emphasis on the evolution of phoresy. Systematic Entomology, 26(1), 33–72. doi: 10.1046/j.1365-3113.2001.00132.x

12. Bolyen, E., Rideout, J. R., Dillon, M. R., Bokulich, N. A., Abnet, C. C., Al-Ghalith, G. A., … Caporaso, J. G. (2019). Reproducible, interactive, scalable and extensible microbiome data science using QIIME 2. Nature Biotechnology, 37(8), 852–857. doi: 10.1038/s41587-019-0209-9

13. Bright, M., & Bulgheresi, S. (2010). A complex journey: transmission of microbial symbionts. Nature Reviews Microbiology, 8(3), 218–230. doi: 10.1038/nrmicro2262

14. Brooks, A. W., Kohl, K. D., Brucker, R. M., Opstal, E. J. van, & Bordenstein, S. R. (2016). Phylosymbiosis: Relationships and Functional Effects of Microbial Communities across Host Evolutionary History. PLOS Biology, 14(11), e2000225. doi: 10.1371/journal.pbio.2000225

15. Callahan, B. J., McMurdie, P. J., Rosen, M. J., Han, A. W., Johnson, A. J. A., & Holmes, S. P. (2016). DADA2: High resolution sample inference from Illumina amplicon data. Nature Methods, 13(7), 581–583. doi: 10.1038/nmeth.3869

16. Cammaerts, R. (1992). Stimuli inducing the regurgitation of the workers of Lasius flavus (Formicidae) upon the myrmecophilous beetle Claviger testaceus (Pselaphidae). Behavioural Processes, 28(1–2), 81–96.

17. Colman, D. R., Toolson, E. C., & Takacs-Vesbach, C. D. (2012). Do diet and taxonomy influence insect gut bacterial communities? Molecular Ecology, 21(20), 5124–5137. doi: 10.1111/j.1365-294x.2012.05752.x

18. Danoff-Burg, J. A. (1994). Evolving under myrmecophily: a cladistic revision of the symphilic beetle tribe Sceptobiini (Coleoptera: Staphylinidae: Aleocharinae). Systematic Entomology, 19(1), 25–45. doi: 10.1111/j.1365-3113.1994.tb00577.x

19. Danoff-Burg, J. A. (1996). An ethogram of the ant-guest beetle tribe Sceptobiini (Coleoptera: Staphylinidae; Formicidae). Sociobiology, 27(3), 287–328.

20. Danoff-Burg, J. A. (2002). Evolutionary Lability and Phylogenetic Utility of Behavior in a Group of Ant-Guest Staphylinidae Beetles. Annals of the Entomological Society of America, 95(2), 143– 155. doi: 10.1603/0013-8746(2002)095[0143:elapuo]2.0.co;2

21. Davis, N. M., Proctor, D. M., Holmes, S. P., Relman, D. A., & Callahan, B. J. (2018). Simple statistical identification and removal of contaminant sequences in marker-gene and meta-genomics data. Microbiome, 6(1), 226. doi: 10.1186/s40168-018-0605-2

22. Dill-McFarland, K. A., Tang, Z.-Z., Kemis, J. H., Kerby, R. L., Chen, G., Palloni, A., … Herd, P. (2019). Close social relationships correlate with human gut microbiota composition. Scientific Reports, 9(1), 703. doi: 10.1038/s41598-018-37298-9

23. Dormann, C. F., Gruber, B., & Fründ, J. (2008). Introducing the bipartite package: analysing ecological networks. Interaction, *1*(0.2413793).

24. Dosmann, A., Bahet, N., & Gordon, D. M. (2016). Experimental modulation of external microbiome affects nestmate recognition in harvester ants (Pogonomyrmex barbatus). PeerJ, 4, e1566. doi: 10.7717/peerj.1566

25. Douglas, A. E. (1998). Nutritional Interactions in Insect-Microbial Symbioses: Aphids and Their Symbiotic Bacteria Buchnera. Annual Review of Entomology, 43(1), 17–37. doi: 10.1146/an-nurev.ento.43.1.17

26. Elmes, G. W., Barr, B., & Thomas, J. A. (1999). Extreme host specificity by Microdon mutabilis (Diptera: Syrphiae), a social parasite of ants. Proceedings of the Royal Society B: Biological Sciences, 266(1418), 447–453. doi: 10.1098/rspb.1999.0658

27. Emms, D. M., & Kelly, S. (2019). OrthoFinder: phylogenetic orthology inference for comparative genomics. Genome Biology, 20(1), 238. doi: 10.1186/s13059-019-1832-y

28. Engel, P., & Moran, N. A. (2013). The gut microbiota of insects – diversity in structure and function. FEMS Microbiology Reviews, 37(5), 699–735. doi: 10.1111/1574-6976.12025

29. Engl, T., Eberl, N., Gorse, C., Krüger, T., Schmidt, T. H. P., Plarre, R., … Kaltenpoth, M. (2018). Ancient symbiosis confers desiccation resistance to stored grain pest beetles. Molecular Ecology, 27(8), 2095–2108. doi: 10.1111/mec.14418

30. Engl, T., & Kaltenpoth, M. (2018). Influence of microbial symbionts on insect pheromones. Natural Product Reports, 35(5), 386–397. doi: 10.1039/c7np00068e

31. Eren, A. M., Esen, Ö. C., Quince, C., Vineis, J. H., Morrison, H. G., Sogin, M. L., & Delmont, T. O. (2015). Anvi’o: an advanced analysis and visualization platform for ‘omics data. PeerJ, 3, e1319. doi: 10.7717/peerj.1319

32. Eren, A. M., Kiefl, E., Shaiber, A., Veseli, I., Miller, S. E., Schechter, M. S., … Willis, A. D. (2021). Community-led, integrated, reproducible multi-omics with anvi’o. Nature Microbiology, 6(1), 3–6. doi: 10.1038/s41564-020-00834-3

33. Feener, D. H., & Brown, B. V. (1997). Diptera as parasitoids. Annual Review of Entomology, 42(1), 73–97. doi: 10.1146/annurev.ento.42.1.73

34. Feldhaar, H., Straka, J., Krischke, M., Berthold, K., Stoll, S., Mueller, M. J., & Gross, R. (2007). Nutritional upgrading for omnivorous carpenter ants by the endosymbiont Blochmannia. BMC Biology, 5(1), 48. doi: 10.1186/1741-7007-5-48

35. Gil, R., Silva, F. J., Zientz, E., Delmotte, F., González-Candelas, F., Latorre, A., … Moya, A. (2003). The genome sequence of Blochmannia floridanus: Comparative analysis of reduced genomes. Proceedings of the National Academy of Sciences, 100(16), 9388–9393. doi: 10.1073/pnas.1533499100

36. Gillespie, J. J., Driscoll, T. P., Verhoeve, V. I., Rahman, M. S., Macaluso, K. R., & Azad, A. F. (2018). A tangled web: origins of reproductive parasitism. Genome Biology and Evolution, 10(9), evy159. doi: 10.1093/gbe/evy159

37. Godfray, H. C. J. (1994). Parasitoids: behavioral and evolutionary ecology (Vol. 67). Princeton University Press.

38. Grimaldi, D. A., & Engel, M. S. (2005). Evolution of the Insects. Cambridge University Press.

39. Hagimori, T., Abe, Y., Date, S., & Miura, K. (2006). The First Finding of a Rickettsia Bacterium Associated with Parthenogenesis Induction Among Insects. Current Microbiology, 52(2), 97–101. doi: 10.1007/s00284-005-0092-0

40. Hawkins, B. A. (1994). Pattern and Process in Host-Parasitoid Interactions. Cambridge University Press. doi: 10.1017/cbo9780511721885

41. Hebard, M. (1920). A Revision of the North American Species of the Genus Myrmecophila (Orthoptera; Gryllidae; Myrmecophilinae). Transactions of the American Entomological Society (1890-), 46(1), 91–111. Retrieved from http://www.jstor.org/stable/25077026

42. Henderson, G., & Akre, R. D. (1986). Biology of the Myrmecophilous Cricket, Myrmecophila manni (Orthoptera: Gryllidae). Journal of the Kansas Entomological Society, 59(3), 454–467. Retrieved from http://www.jstor.org/stable/25084806

43. Hendry, T. A., Hunter, M. S., & Baltrus, D. A. (2014). The Facultative Symbiont Rickettsia Protects an Invasive Whitefly against Entomopathogenic Pseudomonas syringae Strains. Applied and Environmental Microbiology, 80(23), 7161–7168. doi: 10.1128/aem.02447-14

44. Hoang, D. T., Chernomor, O., Haeseler, A. von, Minh, B. Q., & Vinh, L. S. (2017). UFBoot2: Improving the Ultrafast Bootstrap Approximation. Molecular Biology and Evolution, 35(2), 518–522. doi: 10.1093/molbev/msx281

45. Hoey-Chamberlain, R., Rust, M. K., & Klotz, J. H. (2013). A Review of the Biology, Ecology and Behavior of Velvety Tree Ants of North America. Sociobiology, 60(1), 1–10. doi: 10.1155/1914/69251

46. Hölldobler, B. (1967). Zur Physiologie der Gast-Wirt-Beziehungen (Myrmecophilie) bei Ameisen. I. Das Gastverhältnis der Atemeles- und Lomechusa-Larven (Col. Staphylinidae) zu Formica (Hym. Formicidae).*. Zeitschrift für vergleichende Physiologie, 56(1), 1–21. doi: 10.1007/bf00333561

47. Hölldobler, B. (1970). Zur Physiologie der Gast-Wirt-Beziehungen (Myrmecophilie) bei Ameisen. I. Das Gastverhältnis des imaginalen Atemeles pubicollis Bris. (Col. Staphylinidae) zu Myrmica und Formica (Hym. Formicidae). Zeitschrift für vergleichende Physiologie, 66(2), 215– 250. doi: 10.1007/bf00297780

48. Hölldobler, B. (1971). Communication between Ants and Their Guests. Scientific American, 224(3), 86–93. doi: 10.1038/scientificamerican0371-86

49. Hölldobler, B., & Kwapich, C. L. (2019). Behavior and exocrine glands in the myrmecophilous beetle Dinarda dentata (Gravenhorst, 1806) (Coleoptera: Staphylinidae: Aleocharinae). PLOS ONE, 14(1), e0210524–22. doi: 10.1371/journal.pone.0210524

50. Hölldobler, B., & Wilson, E. O. (1990). The Ants. Harvard University Press.

51. Ivens, A. B. F., Gadau, A., Kiers, E. T., & Kronauer, D. J. C. (2018). Can social partnerships influence the microbiome? Insights from ant farmers and their trophobiont mutualists. Molecular Ecology, 27(8), 1898–1914. doi: 10.1111/mec.14506

52. Jennings, E. C., Korthauer, M. W., Hamilton, T. L., & Benoit, J. B. (2019). Matrotrophic viviparity constrains microbiome acquisition during gestation in a live-bearing cockroach, Diploptera punctata. Ecology and Evolution, 9(18), 10601–10614. doi: 10.1002/ece3.5580

53. Jiang, L., Amir, A., Morton, J. T., Heller, R., Arias-Castro, E., & Knight, R. (2017). Discrete False-Discovery Rate Improves Identification of Differentially Abundant Microbes. MSystems, 2(6), e00092–17. doi: 10.1128/msystems.00092-17

54. Jones, P., Binns, D., Chang, H.-Y., Fraser, M., Li, W., McAnulla, C., … Hunter, S. (2014). Inter-ProScan 5: genome-scale protein function classification. Bioinformatics, 30(9), 1236–1240. doi: 10.1093/bioinformatics/btu031

55. Jones, R. T., Sanchez, L. G., & Fierer, N. (2013). A Cross-Taxon Analysis of Insect-Associated Bacterial Diversity. PLoS ONE, 8(4), e61218. doi: 10.1371/journal.pone.0061218

56. Jordan, K. (1913). Zur Morphologie und Biologie der myrmecophilen Gattungen Lomechusa und Atemeles und einiger verwandter Formen. Zeitschrift Für Wissenschaftliche Zoologie, 107, 346–386.

57. Kaczmarczyk-Ziemba, A., Zagaja, M., Wagner, G. K., Pietrykowska-Tudruj, E., & Staniec, B. (2020). First Insight into Microbiome Profiles of Myrmecophilous Beetles and Their Host, Red Wood Ant Formica polyctena (Hymenoptera: Formicidae)—A Case Study. Insects, 11(2), 134. doi: 10.3390/insects11020134

58. Kaltenpoth, M., & Engl, T. (2014). Defensive microbial symbionts in Hymenoptera. Functional Ecology, 28(2), 315–327. doi: 10.1111/1365-2435.12089

59. Kalyaanamoorthy, S., Minh, B. Q., Wong, T. K. F., Haeseler, A. von, & Jermiin, L. S. (2017). ModelFinder: fast model selection for accurate phylogenetic estimates. Nature Methods, 14(6), 587–589. doi: 10.1038/nmeth.4285

60. Kang, D. D., Li, F., Kirton, E., Thomas, A., Egan, R., An, H., & Wang, Z. (2019). MetaBAT 2: an adaptive binning algorithm for robust and efficient genome reconstruction from metagenome assemblies. PeerJ, 7, e7359. doi: 10.7717/peerj.7359

61. Kathirithamby, J. (2009). Host-Parasitoid Associations in Strepsiptera. Annual Review of Entomology, 54(1), 227–249. doi: 10.1146/annurev.ento.54.110807.090525

62. Kistner, D. H. (1979). Social and evolutionary significance of social insect symbionts (H. R. Hermann, Ed.). In Vol. *I* (pp. 339–413).

63. Kistner, D. H. (1982). The Social Insects’ Bestiary (H. R. Hermann, Ed.). In (pp. 1–244).

64. Koch, H., & Schmid-Hempel, P. (2011). Socially transmitted gut microbiota protect bumble bees against an intestinal parasite. Proceedings of the National Academy of Sciences, 108(48), 19288–19292. doi: 10.1073/pnas.1110474108

65. Kolde, R., & Kolde, M. R. (2015). Package ‘pheatmap.’ R Package, 1(7), 790.

66. Komatsu, T., Maruyama, M., & Itino, T. (2009). Behavioral differences between two ant cricket species in Nansei Islands: host-specialist versus host-generalist. Insectes Sociaux, 56(4), 389–396. doi: 10.1007/s00040-009-0036-y

67. Kovarik, P. W., & Caterino, M. S. (2005). Histeridae Gyllenhal, 1808 (R. G. Beutel & R. A. B. Leschen, Eds.). In (pp. 190–222). Walter de Gruyter.

68. Kurtz, S., Phillippy, A., Delcher, A. L., Smoot, M., Shumway, M., Antonescu, C., & Salzberg, S. L. (2004). Versatile and open software for comparing large genomes. Genome Biology, 5(2), R12. doi: 10.1186/gb-2004-5-2-r12

69. Lenoir, A., Chalon, Q., Carvajal, A., Ruel, C., Barroso, Á., Lackner, T., & Boulay, R. (2012). Chemical Integration of Myrmecophilous Guests in Aphaenogaster Ant Nests. Psyche: A Journal of Entomology, 2012, 1–12. doi: 10.1155/2012/840860

70. Leschen, R. A. B. (1991). Behavioral observations on the myrmecophile Fustiger knausii (Coleoptera: Pselaphidae: Clavigerinae) with a discussion of grasping notches in myrmecophiles. Entomological News, 102(5), 215–222.

71. Letunic, I., & Bork, P. (2007). Interactive Tree Of Life (iTOL): an online tool for phylogenetic tree display and annotation. Bioinformatics, 23(1), 127–128. doi: 10.1093/bioinformatics/btl529

72. Li, D., Liu, C.-M., Luo, R., Sadakane, K., & Lam, T.-W. (2015). MEGAHIT: an ultra-fast singlenode solution for large and complex metagenomics assembly via succinct de Bruijn graph. Bioinformatics, 31(10), 1674–1676. doi: 10.1093/bioinformatics/btv033

73. Li, H. (2018). Minimap2: pairwise alignment for nucleotide sequences. Bioinformatics, 34(18), 3094–3100. doi: 10.1093/bioinformatics/bty191

74. Li, H., Handsaker, B., Wysoker, A., Fennell, T., Ruan, J., Homer, N., … Subgroup, 1000 Genome Project Data Processing. (2009). The Sequence Alignment/Map format and SAMtools. Bioinformatics, 25(16), 2078–2079. doi: 10.1093/bioinformatics/btp352

75. López-Estrada, E. K., Sanmartín, I., Uribe, J. E., Abalde, S., & García-París, M. (2021). Diversification dynamics of hypermetamorphic blister beetles (Meloidae): Are homoplastic host shifts and phoresy key factors of a rushing forward strategy to escape extinction? BioRxiv, 2021.01.04.425192. doi: 10.1101/2021.01.04.425192

76. Lozupone, C. A., Hamady, M., Kelley, S. T., & Knight, R. (2007). Quantitative and Qualitative β Diversity Measures Lead to Different Insights into Factors That Structure Microbial Communities. Applied and Environmental Microbiology, 73(5), 1576–1585. doi: 10.1128/aem.01996-06

77. Łukasik, P., Guo, H., Asch, M., Ferrari, J., & Godfray, H. C. J. (2013). Protection against a fungal pathogen conferred by the aphid facultative endosymbionts Rickettsia and Spiroplasma is expressed in multiple host genotypes and species and is not influenced by co-infection with another symbiont. Journal of Evolutionary Biology, 26(12), 2654–2661. doi: 10.1111/jeb.12260

78. Majumder, R., Sutcliffe, B., Adnan, S. M., Mainali, B., Dominiak, B. C., Taylor, P. W., & Chapman, T. A. (2020). Artificial Larval Diet Mediates the Microbiome of Queensland Fruit Fly. Frontiers in Microbiology, 11, 576156. doi: 10.3389/fmicb.2020.576156

79. Maruyama, M., Akino, T., Hashim, R., & Komatsu, T. (2009). Behavior and cuticular hydrocarbons of myrmecophilous insects (Coleoptera: Staphylinidae; Diptera: Phoridae; Thysanura) associated with Asian Aenictus army ants (Hymenoptera; Formicidae). Sociobiology, 54(1), 19–35.

80. Maruyama, M., & Parker, J. (2017). Deep-Time Convergence in Rove Beetle Symbionts of Army Ants. Current Biology, 27(6), 920–926. doi: 10.1016/j.cub.2017.02.030

81. Mason, C. J., Clair, A. St., Peiffer, M., Gomez, E., Jones, A. G., Felton, G. W., & Hoover, K. (2020). Diet influences proliferation and stability of gut bacterial populations in herbivorous lepidopteran larvae. PLOS ONE, 15(3), e0229848. doi: 10.1371/journal.pone.0229848

82. McCutcheon, J. P. (2021). The Genomics and Cell Biology of Host-Beneficial Intracellular Infections. Annual Review of Cell and Developmental Biology, 37(1), 1–28. doi: 10.1146/annurev-cellbio-120219-024122

83. McCutcheon, J. P., & Moran, N. A. (2012). Extreme genome reduction in symbiotic bacteria. Nature Reviews Microbiology, 10(1), 13–26. doi: 10.1038/nrmicro2670

84. McMurdie, P. J., & Holmes, S. (2013). phyloseq: An R Package for Reproducible Interactive Analysis and Graphics of Microbiome Census Data. PLoS ONE, 8(4), e61217. doi: 10.1371/journal.pone.0061217

85. Meer, R. K. V., & Wojcik, D. P. (1982). Chemical Mimicry in the Myrmecophilous Beetle Myrmecaphodius excavaticollis. Science, 218(4574), 806–808. doi: 10.1126/science.218.4574.806

86. Minh, B. Q., Schmidt, H. A., Chernomor, O., Schrempf, D., Woodhams, M. D., Haeseler, A. von, &Lanfear, R. (2020). IQ-TREE 2: New models and efficient methods for phylogenetic inference in the genomic era. Molecular Biology and Evolution, 37(5), 1530–1534. doi: 10.1093/mol-bev/msaa015

87. Moeller, A. H., Foerster, S., Wilson, M. L., Pusey, A. E., Hahn, B. H., & Ochman, H. (2016). Social behavior shapes the chimpanzee pan-microbiome. Science Advances, 2(1), e1500997. doi: 10.1126/sciadv.1500997

88. Moran, N. A., McCutcheon, J. P., & Nakabachi, A. (2008). Genomics and evolution of heritable bacterial symbionts. Annual Review of Genetics, 42(1), 165–190. doi: 10.1146/annurev.genet.41.110306.130119

89. Morton, W., William. (1900). The Habits of Myrmecophila Nebrascensis Bruner. Psyche: A Journal of Entomology, 9(294), 111–115. doi: 10.1155/1900/75323

90. Noh, M. Y., Muthukrishnan, S., Kramer, K. J., & Arakane, Y. (2016). Cuticle formation and pigmentation in beetles. Current Opinion in Insect Science, 17, 1–9. doi: 10.1016/j.cois.2016.05.004

91. Ogle, D., Wheeler, P., & Dinno, A. (2020). FSA: fisheries stock analysis. R package version 0.8. 30. Retrived from https://Github.Com/Droglenc/FSA.

92. Oksanen, J., Blanchet, F. G., Kindt, R., Legendre, P., Minchin, P., O’Hara, R., … others. (2015). Vegan community ecology package: ordination methods, diversity analysis and other functions for community and vegetation ecologists. R Package Ver, 2–3.

93. Oliver, K. M., Russell, J. A., Moran, N. A., & Hunter, M. S. (2003). Facultative bacterial symbionts in aphids confer resistance to parasitic wasps. Proceedings of the National Academy of Sciences, 100(4), 1803–1807. doi: 10.1073/pnas.0335320100

94. Orlov, I., Newton, A. F., & Solodovnikov, A. (2021). Phylogenetic review of the tribal system of Aleocharinae, a mega-lineage of terrestrial arthropods in need of reclassification. Journal of Zoological Systematics and Evolutionary Research. doi: 10.1111/jzs.12524

95. Park, R., Dzialo, M. C., Spaepen, S., Nsabimana, D., Gielens, K., Devriese, H., … Verstrepen, K. J. (2019). Microbial communities of the house fly Musca domestica vary with geographical location and habitat. Microbiome, 7(1), 147. doi: 10.1186/s40168-019-0748-9

96. Parker, J. (2016). Myrmecophily in beetles (Coleoptera): evolutionary patterns and biological mechanisms. Myrmecological News, 22, 65–108.

97. Parker, J., & Grimaldi, D. A. (2014). Specialized Myrmecophily at the Ecological Dawn of Modern Ants. Current Biology, 24(20), 2428–2434. doi: 10.1016/j.cub.2014.08.068

98. Parker, J., & Kronauer, D. J. C. (2021). How ants shape biodiversity. Current Biology.

99. Parks, D. H., Imelfort, M., Skennerton, C. T., Hugenholtz, P., & Tyson, G. W. (2015). CheckM: assessing the quality of microbial genomes recovered from isolates, single cells, and meta-genomes. Genome Research, 25(7), 1043–1055. doi: 10.1101/gr.186072.114

100. Perlman, S. J., Hunter, M. S., & Zchori-Fein, E. (2006). The emerging diversity of Rickettsia. Proceedings of the Royal Society B: Biological Sciences, 273(1598), 2097–2106. doi: 10.1098/rspb.2006.3541

101. Perotti, M. A., Clarke, H. K., Turner, B. D., & Braig, H. R. (2006). Rickettsia as obligate and mycetomic. The FASEB Journal, 20(13), 2372–2374. doi: 10.1096/fj.06-5870fje

102. Piel, J. (2002). A polyketide synthase-peptide synthetase gene cluster from an uncultured bacterial symbiont of Paederus beetles. Proceedings of the National Academy of Sciences of the United States of America, 99(22), 14002–14007. doi: 10.1073/pnas.222481399

103. Pierce, N. E., Braby, M. F., Heath, A., Lohman, D. J., Mathew, J., Rand, D. B., & Travassos, M. A. (2002). The ecology and evolution of ant association in the Lycaenidae (Lepidoptera). Annual Review of Entomology, 47, 733–771. doi: 10.1146/annurev.ento.47.091201.145257

104. Quast, C., Pruesse, E., Yilmaz, P., Gerken, J., Schweer, T., Yarza, P., … Glöckner, F. O. (2013). The SILVA ribosomal RNA gene database project: improved data processing and web-based tools. Nucleic Acids Research, 41(D1), D590–D596. doi: 10.1093/nar/gks1219

105. Renelies-Hamilton, J., Germer, K., Sillam-Dussès, D., Bodawatta, K. H., & Poulsen, M. (2021). Disentangling the Relative Roles of Vertical Transmission, Subsequent Colonizations, and Diet on Cockroach Microbiome Assembly. MSphere, 6(1). doi: 10.1128/msphere.01023-20

106. Russell, J. A., Moreau, C. S., Goldman-Huertas, B., Fujiwara, M., Lohman, D. J., & Pierce, N. E. (2009). Bacterial gut symbionts are tightly linked with the evolution of herbivory in ants. Proceedings of the National Academy of Sciences, 106(50), 21236–21241. doi: 10.1073/pnas.0907926106

107. Salem, H., Bauer, E., Kirsch, R., Berasategui, A., Cripps, M., Weiss, B., … Kaltenpoth, M. (2017). Drastic Genome Reduction in an Herbivore’s Pectinolytic Symbiont. Cell, 1–26. doi: 10.1016/j.cell.2017.10.029

108. Salter, S. J., Cox, M. J., Turek, E. M., Calus, S. T., Cookson, W. O., Moffatt, M. F., … Walker, A.W. (2014). Reagent and laboratory contamination can critically impact sequence-based microbiome analyses. BMC Biology, 12(1), 87. doi: 10.1186/s12915-014-0087-z

109. Scudder, G. G. E. (2017). The Importance of Insects. In R. G. Foottit & P. H. Adler (Eds.), Insect Biodiversity: Science and Society (pp. 9–43). John Wiley & Sons. doi: 10.1002/9781118945568.ch2

110. Seemann, T. (2014). Prokka: rapid prokaryotic genome annotation. Bioinformatics, 30(14), 2068– 2069. doi: 10.1093/bioinformatics/btu153

111. Seevers, C. H. (1965). The systematics, evolution and zoogeography of staphylinid beetles associated with army ants (Coleoptera, Staphylinidae). Fieldiana Zoology, 47(2), 139–351.

112. Song, S. J., Lauber, C., Costello, E. K., Lozupone, C. A., Humphrey, G., Berg-Lyons, D., … Knight, R. (2013). Cohabiting family members share microbiota with one another and with their dogs. ELife, 2, e00458. doi: 10.7554/elife.00458

113. Stadler, B., & Dixon, A. F. G. (2005). Ecology and Evolution of Aphid-Ant Interactions. Annual Review of Ecology, Evolution, and Systematics, 36, 345–372.

114. Stoeffler, M., Tolasch, T., & Steidle, J. L. M. (2011). Three beetles—three concepts. Different defensive strategies of congeneric myrmecophilous beetles. Behavioral Ecology and Sociobiology, 65(8), 1605–1613. doi: 10.1007/s00265-011-1171-9

115. Strand, M. R., & Obrycki, J. J. (1996). Host Specificity of Insect Parasitoids and PredatorsMany factors influence the host ranges of insect natural enemies. BioScience, 46(6), 422–429. doi: 10.2307/1312876

116. Thayer, M. K. (2005). Staphylinidae Latreille, 1802 (R. G. Beutel & R. A. B. Leschen, Eds.). In (pp. 296–344). Walter de Gruyter.

117. Tung, J., Barreiro, L. B., Burns, M. B., Grenier, J.-C., Lynch, J., Grieneisen, L. E., … Archie, E. A. (2015). Social networks predict gut microbiome composition in wild baboons. ELife, 4, e05224. doi: 10.7554/elife.05224

118. Ul-Hasan, S., Bowers, R. M., Figueroa-Montiel, A., Licea-Navarro, A. F., Beman, J. M., Woyke, T., & Nobile, C. J. (2019). Community ecology across bacteria, archaea and microbial eukaryotes in the sediment and seawater of coastal Puerto Nuevo, Baja California. PLOS ONE, 14(2), e0212355. doi: 10.1371/journal.pone.0212355

119. Vigneron, A., Masson, F., Vallier, A., Balmand, S., Rey, M., Vincent-Monégat, C., … Heddi, A. (2014). Insects Recycle Endosymbionts when the Benefit Is Over. Current Biology, 24(19), 2267–2273. doi: 10.1016/j.cub.2014.07.065

120. Wada-Katsumata, A., Zurek, L., Nalyanya, G., Roelofs, W. L., Zhang, A., & Schal, C. (2015). Gut bacteria mediate aggregation in the German cockroach. Proceedings of the National Academy of Sciences, 112(51), 15678–15683. doi: 10.1073/pnas.1504031112

121. Wang, T. B., Patel, A., Vu, F., & Nonacs, P. (2010). Natural History Observations on the Velvety Tree Ant (Liometopum occidentale): Unicoloniality and Mating Flights. Sociobiology, 55(3), 787–794. doi: 10.1111/j.1558-5646.2009.00628.x

122. Wick, R. R., Judd, L. M., Gorrie, C. L., & Holt, K. E. (2017). Unicycler: Resolving bacterial genome assemblies from short and long sequencing reads. PLOS Computational Biology, 13(6), e1005595. doi: 10.1371/journal.pcbi.1005595

123. Wickham, H. (2011). ggplot2. Wiley Interdisciplinary Reviews: Computational Statistics, 3(2), 180– 185.

124. Wojcik, D. P. (1989). Behavioral Interactions between Ants and Their Parasites. The Florida Entomologist, 72(1), 43–51. doi: 10.2307/3494966?ref=no-x-route:2d46a42242f93ba50fee4895d3959302

125. Yamamoto, S., Maruyama, M., & Parker, J. (2016). Evidence for social parasitism of early insect societies by Cretaceous rove beetles. Nature Communications, 7, 13658. doi: 10.1038/ncomms13658

126. Yun, J.-H., Roh, S. W., Whon, T. W., Jung, M.-J., Kim, M.-S., Park, D.-S., … Bae, J.-W. (2014). Insect Gut Bacterial Diversity Determined by Environmental Habitat, Diet, Developmental Stage, and Phylogeny of Host. Applied and Environmental Microbiology, 80(17), 5254–5264. doi: 10.1128/aem.01226-14

127. Yusuf, M., & Turner, B. (2004). Characterisation of Wolbachia-like bacteria isolated from the parthenogenetic stored-product pest psocid Liposcelis bostrychophila (Badonnel) (Psocoptera). Journal of Stored Products Research, 40(2), 207–225. doi: 10.1016/s0022-474x(02)00098-x

128. Zientz, E., Dandekar, T., & Gross, R. (2004). Metabolic Interdependence of Obligate Intracellular Bacteria and Their Insect Hosts†. Microbiology and Molecular Biology Reviews, 68(4), 745– 770. doi: 10.1128/mmbr.68.4.745-770.2004

