## Supplementary material for "Structure of an ant-myrmecophile-microbe community": Table S2

| Figure 2 | Comparison | N number | Methods | *P* value | Statistical values |
|---|---|---|---|---|---|
| 2E | Effect of species on overall microbiota composition assessed using Jaccard distances | 35 *Liometopum*, 9 *Myrmecophilus*, 44 *Sceptobius*, 2 *Liometoxenus*, 19 *Platyusa*, 4 *Pella*, 5 *Lasius*, 4 *Drusilla*, 5 *Lissagria* | PERMANOVA | 0.001 | $F$ = 3.1348, $R^2$ = 0.1943 |
| | Effect of species on overall microbiota composition assessed using Bray-Curtis dissimilarity | | | 0.001 | $F$ = 14.535, $R^2$ = 0.5279 |
| 2F | Effect of colony identity (nest site) on overall microbiota composition assessed using Jaccard distances | 33 *Liometopum*, 9 *Myrmecophilus*, 44 *Sceptobius*, 2 *Liometoxenus*, 19 *Platyusa* | PERMANOVA | 0.001 | $F$ = 1.4603, $R^2$ = 0.0490 |
| | Effect of the interaction between species and nest site on overall microbiota composition assessed using Jaccard distances | | | 0.008 | $F$ = 1.1102, $R^2$ = 0.0838 |

| | | | | | |
|---|---|---|---|---|---|
| (not shown) | Effect of colony identity (nest site) on overall microbiota composition assessed using Jaccard distances, restricted to gut samples<br><br>Effect of colony identity (nest site) on overall microbiota composition assessed using Jaccard distances, restricted to head samples | 13 *Liometopum*, 11 *Sceptobius*, 6 *Platyusa*<br><br>14 *Sceptobius*, 5 *Platyusa* | | 0.084<br><br>0.064 | $F$ = 1.1502, $R^2$ = 0.1329<br><br>$F$ = 1.1020, $R^2$ = 0.2258 |
| **Figure 3** | **Comparison** | **N number** | **Methods** | ***P* value** | **Statistical values** |
| 3A | Jaccard distances between *Liometopum* and *Sceptobius*, versus *Liometopum* and *Myrmecophilus*, versus *Liometopum* and *Platyusa* | 144 *Liometopum* vs. *Sceptobius* distances, 66 *Liometopum* vs. *Myrmecophilus* distances, 85 *Liometopum* vs. *Platyusa* distances (derived from 31 *Liometopum* samples, 28 *Sceptobius* samples, 9 *Myrmecophilus* samples, and 13 *Platyusa* samples) | Kruskal-Wallis test | $< 2.2 \times 10^{-16}$ | H = 103.31 |
| | Jaccard distances between *Liometopum* and *Sceptobius*, versus *Liometopum* and *Myrmecophilus* | 144 *Liometopum* vs. *Sceptobius* distances, 66 *Liometopum* vs. *Myrmecophilus* distances | Post-hoc: Dunn's test (*p*-values adjusted with the Holm method) | 0.0018 | Z = 3.1268 |

| | | | | | |
|---|---|---|---|---|---|
| | Jaccard distances between *Liometopum* and *Sceptobius*, versus *Liometopum* and *Platyusa* | 144 *Liometopum* vs. *Sceptobius* distances, 85 *Liometopum* vs. *Platyusa* distances | | $9.1980 \times 10^{-24}$ | Z = 10.1576 |
| | Jaccard distances between *Liometopum* and *Myrmecophilus*, versus *Liometopum* and *Platyusa* | 66 *Liometopum* vs. *Myrmecophilus* distances, 85 *Liometopum* vs. *Platyusa* distances | | $3.48 \times 10^{-8}$ | Z = -5.6356 |
| 3B | Jaccard distances between wash samples of *Liometopum* and *Sceptobius*, versus wash samples of *Liometopum* and *Myrmecophilus*, versus wash samples of *Liometopum* and *Platyusa* | 71 *Liometopum* vs. *Sceptobius* distances, 31 *Liometopum* vs. *Myrmecophilus* distances, 80 *Liometopum* vs. *Platyusa* distances (derived from 23 *Liometopum* samples, 17 *Sceptobius* samples, 9 *Myrmecophilus* samples, and 16 *Platyusa* samples) | Kruskal-Wallis test | $3.084 \times 10^{-9}$ | H = 39.194 |
| | Jaccard distances between wash samples of *Liometopum* and *Sceptobius*, versus wash samples of *Liometopum* and *Myrmecophilus* | 71 *Liometopum* vs. *Sceptobius* distances, 31 *Liometopum* vs. *Myrmecophilus* distances | Post-hoc: Dunn's test (*p*-values adjusted with the Holm method) | 0.8918 | Z = 0.2278 |
| | Jaccard distances between wash samples of *Liometopum* and *Sceptobius*, versus wash samples of *Liometopum* and *Platyusa* | 71 *Liometopum* vs. *Sceptobius* distances, 80 *Liometopum* vs. *Platyusa* distances | | $1.7443 \times 10^{-8}$ | Z = 5.8220 |
| | Jaccard distances between wash samples of *Liometopum* and *Myrmecophilus*, versus wash samples of *Liometopum* and *Platyusa* | 31 *Liometopum* vs. *Myrmecophilus* distances, 80 *Liometopum* vs. *Platyusa* distances | | $4.1785 \times 10^{-5}$ | Z = -4.2551 |

| | | | | | |
|---|---|---|---|---|---|
| 3D | Jaccard distances between *Liometopum* bodies and *Liometopum* guts, versus *Liometopum* bodies and *Sceptobius* heads | 228 *Liometopum* bodies vs. *Liometopum* guts distances, 228 *Liometopum* bodies vs. *Sceptobius* heads distances (derived from 19 *Liometopum* bodies, 12 *Liometopum* guts, and 12 *Sceptobius* heads) | Wilcoxon test with the Benjamini-Hochberg method for controlling the false discovery rate | $7.4405 \times 10^{-6}$ | W = 19482 |
| | Jaccard distances between *Liometopum* bodies and *Sceptobius* heads, versus *Liometopum* bodies and *Sceptobius* bodies | 228 *Liometopum* bodies vs. *Sceptobius* heads distances, 171 *Liometopum* bodies vs. *Sceptobius* bodies (derived from 19 *Liometopum* bodies, 12 *Sceptobius* heads, and 9 *Sceptobius* bodies) | | 0.0090 | W = 16359 |
| | Jaccard distances between *Liometopum* bodies and *Sceptobius* bodies, versus *Liometopum* bodies and *Sceptobius* guts | 171 *Liometopum* bodies vs. *Sceptobius* bodies, 76 *Liometopum* bodies vs. *Sceptobius* guts (derived from 19 *Liometopum* bodies, 9 *Sceptobius* bodies, and 4 *Sceptobius* guts) | | $5.1776 \times 10^{-10}$ | W = 3134.5 |
| | Jaccard distances between *Liometopum* bodies and *Sceptobius* guts, versus *Liometopum* bodies and *Platyusa* bodies | 76 *Liometopum* bodies vs. *Sceptobius* guts, 95 *Liometopum* bodies vs. *Platyusa* bodies (derived from 19 *Liometopum* bodies, 4 *Sceptobius* guts, and 5 *Platyusa* bodies) | | $1.4556 \times 10^{-6}$ | W = 1991 |

| | | | | | |
|---|---|---|---|---|---|
| | Jaccard distances between *Liometopum* bodies and *Platyusa* bodies, versus *Liometopum* bodies and *Platyusa* guts | 95 *Liometopum* bodies vs. *Platyusa* bodies, 95 *Liometopum* bodies vs. *Platyusa* guts (derived from 19 *Liometopum* bodies, 5 *Platyusa* bodies, and 5 *Platyusa* guts) | | 0.9295 | W = 4478.5 |
| | Jaccard distances between *Liometopum* bodies and *Platyusa* guts, versus *Liometopum* bodies and *Platyusa* heads | 95 *Liometopum* bodies vs. *Platyusa* guts, 38 *Liometopum* bodies vs. *Platyusa* heads (derived from 19 *Liometopum* bodies, 5 *Platyusa* guts, and 2 *Platyusa* heads) | | 0.5768 | W = 1664 |
| **Figure S2** | **Comparison** | **N number** | **Methods** | ***P* value** | **Statistical values** |
| S2A | Effect of location on nest microbiota composition, assessed using Jaccard distances | 3 First Tree, 3 Mile High, 3 Post, 3 Rocks, 3 Sleepy Tree | PERMANOVA | 0.003 | $F$ = 1.2803, $R^2$ = 0.3387 |
| | Effect of location on nest microbiota composition, assessed using Bray-Curtis dissimilarity | 3 First Tree, 3 Mile High, 3 Post, 3 Rocks, 3 Sleepy Tree | | 0.039 | $F$ = 1.7922, $R^2$ = 0.4176 |
| S2B | Jaccard distances between nest and wash samples, vs. Jaccard distances between nest and body part samples | 975 Nest vs. Wash distances, 1575 Nest vs. Body Parts distances (derived from 15 Nest samples, 65 Wash samples, and 105 Body Parts samples) | Wilcoxon test | $< 2.2 \times 10^{-16}$ | W= 229314 |

| | | | | | |
|---|---|---|---|---|---|
| | Bray-Curtis dissimilarities between nest and wash samples, vs. Bray-Curtis dissimilarities between nest and body part samples | 975 Nest vs. Wash distances, 1575 Nest vs. Body Parts distances (derived from 15 Nest samples, 65 Wash samples, and 105 Body Parts samples) | | $< 2.2 \times 10^{-16}$ | W = 230802 |
| **Figure S3** | **Comparison** | **N number** | **Methods** | ***P* value** | **Statistical values** |
| S3 | Bray-Curtis dissimilarities between *Liometopum* and *Sceptobius*, versus *Liometopum* and *Myrmecophilus*, versus *Liometopum* and *Platyusa* | 144 *Liometopum* vs. *Sceptobius* dissimilarities, 66 *Liometopum* vs. *Myrmecophilus* dissimilarities, 85 *Liometopum* vs. *Platyusa* dissimilarities (derived from 31 *Liometopum* samples, 28 *Sceptobius* samples, 9 *Myrmecophilus* samples, and 13 *Platyusa* samples) | Kruskal-Wallis test | $5.197 \times 10^{-8}$ | H = 33.545 |
| | Bray-Curtis dissimilarities between *Liometopum* and *Sceptobius*, versus *Liometopum* and *Myrmecophilus* | 144 *Liometopum* vs. *Sceptobius* dissimilarities, 66 *Liometopum* vs. *Myrmecophilus* dissimilarities | Post-hoc: Dunn's test (*p*-values adjusted with the Holm method) | 0.0136 | Z = 2.7062 |
| | Bray-Curtis dissimilarities between *Liometopum* and *Sceptobius*, versus *Liometopum* and *Platyusa* | 144 *Liometopum* vs. *Sceptobius* dissimilarities, 85 *Liometopum* vs. *Platyusa* dissimilarities | | $2.890 \times 10^{-8}$ | Z = 5.7371 |
| | Bray-Curtis dissimilarities between *Liometopum* and *Myrmecophilus*, versus *Liometopum* and *Platyusa* | 66 *Liometopum* vs. *Myrmecophilus* dissimilarities, 85 *Liometopum* vs. *Platyusa* dissimilarities | | 0.0197 | Z = -2.3311 |

| **Figure S4** | **Comparison** | **N number** | **Methods** | ***P* value** | **Statistical values** |
|---|---|---|---|---|---|
| S4A | Bray-Curtis dissimilarities between *Liometopum* bodies and *Liometopum* guts, versus *Liometopum* bodies and *Sceptobius* bodies | 228 *Liometopum* bodies vs. *Liometopum* guts dissimilarities, 171 *Liometopum* bodies vs. *Sceptobius* bodies dissimilarities (derived from 19 *Liometopum* bodies, 12 *Liometopum* guts, and 9 *Sceptobius* bodies) | Wilcoxon test with the Benjamini-Hochberg method for controlling the false discovery rate | 0.0204 | W = 16566 |
| | Bray-Curtis dissimilarities between *Liometopum* bodies and *Sceptobius* bodies, versus *Liometopum* bodies and *Sceptobius* heads | 171 *Liometopum* bodies vs. *Sceptobius* bodies dissimilarities, 228 *Liometopum* bodies vs. *Sceptobius* heads dissimilarities (derived from 19 *Liometopum* bodies, 9 *Sceptobius* bodies, and 12 *Sceptobius* heads) | | 0.8939 | W = 19646 |
| | Bray-Curtis dissimilarities between *Liometopum* bodies and *Sceptobius* heads, versus *Liometopum* bodies and *Sceptobius* guts | 228 *Liometopum* bodies vs. *Sceptobius* heads dissimilarities, 76 *Liometopum* bodies vs. *Sceptobius* guts dissimilarities (derived from 19 *Liometopum* bodies, 12 *Sceptobius* heads, and 4 *Sceptobius* guts) | | $4.928 \times 10^{-5}$ | W = 5804.5 |
| | Bray-Curtis dissimilarities between *Liometopum* bodies and *Sceptobius* guts, versus *Liometopum* bodies and *Platyusa* bodies | 76 *Liometopum* bodies vs. *Sceptobius* guts dissimilarities, 95 *Liometopum* bodies vs. *Platyusa* bodies dissimilarities (derived from 19 *Liometopum* bodies, 4 *Sceptobius* guts, and 5 *Platyusa* bodies) | | $4.928 \times 10^{-5}$ | W = 2213 |

| | | | | | |
|---|---|---|---|---|---|
| | Bray-Curtis dissimilarities between *Liometopum* bodies and *Platyusa* bodies, versus *Liometopum* bodies and *Platyusa* guts | 95 *Liometopum* bodies vs. *Platyusa* bodies dissimilarities, 95 *Liometopum* bodies vs. *Platyusa* guts dissimilarities (derived from 19 *Liometopum* bodies, 5 *Platyusa* bodies, and 5 *Platyusa* guts) | | 0.5529 | W = 4233 |
| | Bray-Curtis dissimilarities between *Liometopum* bodies and *Platyusa* guts, versus *Liometopum* bodies and *Platyusa* heads | 95 *Liometopum* bodies vs. *Platyusa* guts dissimilarities, 38 *Liometopum* bodies vs. *Platyusa* heads dissimilarities (derived from 19 *Liometopum* bodies, 5 *Platyusa* guts, and 2 *Platyusa* heads) | | 0.1767 | W = 1493 |
| S4B | Bray-Curtis dissimilarities between *Liometopum* guts and *Sceptobius* guts, versus *Liometopum* guts and *Liometopum* bodies | 48 *Liometopum* guts vs. *Sceptobius* guts dissimilarities, 228 *Liometopum* guts vs. *Liometopum* bodies dissimilarities (derived from 12 *Liometopum* guts, 4 *Sceptobius* guts, and 19 *Liometopum* bodies) | Wilcoxon test with the Benjamini-Hochberg method for controlling the false discovery rate | 0.8311 | W = 5596.5 |
| | Bray-Curtis dissimilarities between *Liometopum* guts and *Liometopum* bodies, versus *Liometopum* guts and *Sceptobius* heads | 228 *Liometopum* guts vs. *Liometopum* bodies dissimilarities, 144 *Liometopum* guts vs. *Sceptobius* heads dissimilarities (derived from 12 *Liometopum* guts, 19 *Liometopum* bodies, and 12 *Sceptobius* heads) | | 0.8311 | W = 16200 |

| | | | | | |
|---|---|---|---|---|---|
| | Bray-Curtis dissimilarities between *Liometopum* guts and *Sceptobius* heads, versus *Liometopum* guts and *Sceptobius* bodies | 144 *Liometopum* guts vs. *Sceptobius* heads dissimilarities, 108 *Liometopum* guts vs. *Sceptobius* bodies dissimilarities (derived from 12 *Liometopum* guts, 12 *Sceptobius* heads, and 9 *Sceptobius* bodies) | | 0.8311 | W = 7368.5 |
| | Bray-Curtis dissimilarities between *Liometopum* guts and *Sceptobius* bodies, versus *Liometopum* guts and *Platyusa* bodies | 108 *Liometopum* guts vs. *Sceptobius* bodies dissimilarities, 60 *Liometopum* guts vs. *Platyusa* bodies dissimilarities (derived from 12 *Liometopum* guts, 9 *Sceptobius* bodies, and 5 *Platyusa* bodies) | | $2.1613 \times 10^{-16}$ | W = 695 |
| | Bray-Curtis dissimilarities between *Liometopum* guts and *Platyusa* bodies, versus *Liometopum* guts and *Platyusa* guts | 60 *Liometopum* guts vs. *Platyusa* bodies dissimilarities, 60 *Liometopum* guts vs. *Platyusa* guts dissimilarities (derived from 12 *Liometopum* guts, 5 *Platyusa* bodies, and 5 *Platyusa* guts) | | 0.4148 | W = 1518 |
| | Bray-Curtis dissimilarities between *Liometopum* guts and *Platyusa* guts, versus *Liometopum* guts and *Platyusa* heads | 60 *Liometopum* guts vs. *Platyusa* guts dissimilarities, 24 *Liometopum* guts vs. *Platyusa* guts dissimilarities (derived from 12 *Liometopum* guts, 5 *Platyusa* guts, and 2 *Platyusa* heads) | | 0.8311 | W = 679.5 |

| | | | | | |
|---|---|---|---|---|---|
| S4C | Jaccard distances between *Liometopum* guts and *Liometopum* bodies, versus *Liometopum* guts and *Sceptobius* heads | 228 *Liometopum* guts vs. *Liometopum* bodies distances, 144 *Liometopum* guts vs. *Sceptobius* heads distances (derived from 12 *Liometopum* guts, 19 *Liometopum* bodies, and 12 *Sceptobius* heads) | Wilcoxon test with the Benjamini-Hochberg method for controlling the false discovery rate | $6.8820 \times 10^{-10}$ | W = 9903.5 |
| | Jaccard distances between *Liometopum* guts and *Sceptobius* heads, versus *Liometopum* guts and *Sceptobius* bodies | 144 *Liometopum* guts vs. *Sceptobius* heads distances, 108 *Liometopum* guts vs. *Sceptobius* bodies distances (derived from 12 *Liometopum* guts, 12 *Sceptobius* heads, and 9 *Sceptobius* bodies) | | 0.3696 | W = 7113.5 |
| | Jaccard distances between *Liometopum* guts and *Sceptobius* bodies, versus *Liometopum* guts and *Sceptobius* guts | 108 *Liometopum* guts vs. *Sceptobius* bodies distances, 48 *Liometopum* guts vs. *Sceptobius* guts distances (derived from 12 *Liometopum* guts, 9 *Sceptobius* bodies, and 4 *Sceptobius* guts) | | 0.3696 | W = 2326 |
| | Jaccard distances between *Liometopum* guts and *Sceptobius* guts, versus *Liometopum* guts and *Platyusa* guts | 48 *Liometopum* guts vs. *Sceptobius* guts distances, 60 *Liometopum* guts vs. *Platyusa* guts distances (derived from 12 *Liometopum* guts, 4 *Sceptobius* guts, and 5 *Platyusa* guts) | | $3.9620 \times 10^{-6}$ | W = 658.5 |
| | Jaccard distances between *Liometopum* guts and *Platyusa* guts, versus *Liometopum* guts and *Platyusa* bodies | 60 *Liometopum* guts vs. *Platyusa* guts distances, 60 *Liometopum* guts vs. *Platyusa* bodies distances (derived from 12 *Liometopum* guts, 5 *Platyusa* guts, and 5 *Platyusa* bodies) | | 0.7050 | W = 1872.5 |

| | | | | | |
|---|---|---|---|---|---|
| | Jaccard distances between *Liometopum* guts and *Platyusa* bodies, versus *Liometopum* guts and *Platyusa* heads | 60 *Liometopum* guts vs. *Platyusa* bodies distances, 24 *Liometopum* guts vs. *Platyusa* heads distances (derived from 12 *Liometopum* guts, 5 *Platyusa* bodies, and 2 *Platyusa* heads) | | 0.3696 | W = 599.5 |
| S4D | Bray-Curtis dissimilarities between *Liometopum* wash samples and *Liometopum* bodies, versus *Liometopum* wash samples and *Sceptobius* heads | 437 *Liometopum* wash samples vs. *Liometopum* bodies dissimilarities, 276 *Liometopum* wash samples vs. *Sceptobius* heads dissimilarities (derived from 23 *Liometopum* wash samples, 19 *Liometopum* bodies, and 12 *Sceptobius* heads) | Wilcoxon test with the Benjamini-Hochberg method for controlling the false discovery rate | $5.9037 \times 10^{-31}$ | W = 28906 |
| | Bray-Curtis dissimilarities between *Liometopum* wash samples and *Sceptobius* heads, versus *Liometopum* wash samples and *Sceptobius* bodies | 276 *Liometopum* wash samples vs. *Sceptobius* heads dissimilarities, 207 *Liometopum* wash samples vs. *Sceptobius* bodies dissimilarities (derived from 23 *Liometopum* wash samples, 12 *Sceptobius* heads, and 9 *Sceptobius* bodies) | | 0.2251 | W = 26724 |
| | Bray-Curtis dissimilarities between *Liometopum* wash samples and *Sceptobius* bodies, versus *Liometopum* wash samples and *Sceptobius* guts | 207 *Liometopum* wash samples vs. *Sceptobius* bodies dissimilarities, 92 *Liometopum* wash samples vs. *Sceptobius* guts dissimilarities (derived from 23 *Liometopum* wash samples, 9 *Sceptobius* bodies, and 4 *Sceptobius* guts) | | 0.0002 | W = 6816.5 |

| | | | | | |
|---|---|---|---|---|---|
| | Bray-Curtis dissimilarities between *Liometopum* wash samples and *Sceptobius* guts, versus *Liometopum* wash samples and *Platyusa* bodies | 92 *Liometopum* wash samples vs. *Sceptobius* guts dissimilarities, 115 *Liometopum* wash samples vs. *Platyusa* bodies dissimilarities (derived from 23 *Liometopum* wash samples, 4 *Sceptobius* guts, and 5 *Platyusa* bodies) | | $1.3758 \times 10^{-9}$ | W = 2623 |
| | Bray-Curtis dissimilarities between *Liometopum* wash samples and *Platyusa* bodies, versus *Liometopum* wash samples and *Platyusa* guts | 115 *Liometopum* wash samples vs. *Platyusa* bodies dissimilarities, 115 *Liometopum* wash samples vs. *Platyusa* guts dissimilarities (derived from 23 *Liometopum* wash samples, 5 *Platyusa* bodies, and 5 *Platyusa* guts) | | 0.1056 | W = 5701 |
| | Bray-Curtis dissimilarities between *Liometopum* wash samples and *Platyusa* guts, versus *Liometopum* wash samples and *Platyusa* heads | 115 *Liometopum* wash samples vs. *Platyusa* guts dissimilarities, 46 *Liometopum* wash samples vs. *Platyusa* guts dissimilarities (derived from 23 *Liometopum* wash samples, 5 *Platyusa* guts, and 2 *Platyusa* heads) | | 0.2179 | W = 2290.5 |
| S4E | Weighted UniFrac distances between *Liometopum* bodies and *Liometopum* guts, versus *Liometopum* bodies and *Sceptobius* heads | 228 *Liometopum* bodies vs. *Liometopum* guts distances, 228 *Liometopum* bodies vs. *Sceptobius* heads distances (derived from 19 *Liometopum* bodies, 12 *Liometopum* guts, and 12 *Sceptobius* heads) | Wilcoxon test with the Benjamini-Hochberg method for controlling the false discovery rate | 0.7201 | W = 25296 |

| | | | | | |
|---|---|---|---|---|---|
| | Weighted UniFrac distances between *Liometopum* bodies and *Liometopum* guts, versus *Liometopum* bodies and *Sceptobius* guts | 228 *Liometopum* bodies vs. *Liometopum* guts distances, 76 *Liometopum* bodies vs. *Sceptobius* guts distances (derived from 19 *Liometopum* bodies, 12 *Liometopum* guts, and 4 *Sceptobius* guts) | | 0.5305 | W = 9280 |
| | Weighted UniFrac distances between *Liometopum* bodies and *Liometopum* guts, versus *Liometopum* bodies and *Sceptobius* bodies | 228 *Liometopum* bodies vs. *Liometopum* guts distances, 171 *Liometopum* bodies vs. *Sceptobius* bodies distances (derived from 19 *Liometopum* bodies, 12 *Liometopum* guts, and 9 *Sceptobius* bodies) | | 0.7201 | W = 19085 |
| | Weighted UniFrac distances between *Liometopum* bodies and *Liometopum* guts, versus *Liometopum* bodies and *Platyusa* bodies | 228 *Liometopum* bodies vs. *Liometopum* guts distances, 95 *Liometopum* bodies vs. *Platyusa* bodies distances (derived from 19 *Liometopum* bodies, 12 *Liometopum* guts, and 5 *Platyusa* bodies) | | $5.2830 \times 10^{-8}$ | W = 6521 |
| | Weighted UniFrac distances between *Liometopum* bodies and *Liometopum* guts, versus *Liometopum* bodies and *Platyusa* heads | 228 *Liometopum* bodies vs. *Liometopum* guts distances, 38 *Liometopum* bodies vs. *Platyusa* heads distances (derived from 19 *Liometopum* bodies, 12 *Liometopum* guts, and 2 *Platyusa* heads) | | $7.4529 \times 10^{-4}$ | W = 2769 |

| | | | | | |
|---|---|---|---|---|---|
| | Weighted UniFrac distances between *Liometopum* bodies and *Liometopum* guts, versus *Liometopum* bodies and *Platyusa* guts | 228 *Liometopum* bodies vs. *Liometopum* guts distances, 95 *Liometopum* bodies vs. *Platyusa* guts distances (derived from 19 *Liometopum* bodies, 12 *Liometopum* guts, and 5 *Platyusa* guts) | | $3.2206 \times 10^{-8}$ | W = 6367 |
| S4F | Weighted UniFrac distances between *Liometopum* guts and *Liometopum* bodies, versus *Liometopum* guts and *Sceptobius* heads | 228 *Liometopum* guts vs. *Liometopum* bodies distances, 144 *Liometopum* guts vs. *Sceptobius* heads distances (derived from 12 *Liometopum* guts, 19 *Liometopum* bodies, and 12 *Sceptobius* heads) | Wilcoxon test with the Benjamini-Hochberg method for controlling the false discovery rate | $9.4879 \times 10^{-4}$ | W = 12885 |
| | Weighted UniFrac distances between *Liometopum* guts and *Liometopum* bodies, versus *Liometopum* guts and *Sceptobius* bodies | 228 *Liometopum* guts vs. *Liometopum* bodies distances, 108 *Liometopum* guts vs. *Sceptobius* bodies distances (derived from 12 *Liometopum* guts, 19 *Liometopum* bodies, and 9 *Sceptobius* bodies) | | 0.0856 | W = 10812 |
| | Weighted UniFrac distances between *Liometopum* guts and *Liometopum* bodies, versus *Liometopum* guts and *Sceptobius* guts | 228 *Liometopum* guts vs. *Liometopum* bodies distances, 48 *Liometopum* guts vs. *Sceptobius* guts distances (derived from 12 *Liometopum* guts, 19 *Liometopum* bodies, and 4 *Sceptobius* guts) | | 0.1273 | W = 4705 |

| | | | | | |
|---|---|---|---|---|---|
| | Weighted UniFrac distances between *Liometopum* guts and *Liometopum* bodies, versus *Liometopum* guts and *Platyusa* heads | 228 *Liometopum* guts vs. *Liometopum* bodies distances, 24 *Liometopum* guts vs. *Platyusa* heads distances (derived from 12 *Liometopum* guts, 19 *Liometopum* bodies, and 2 *Platyusa* heads) | | 0.0476 | W = 2006 |
| | Weighted UniFrac distances between *Liometopum* guts and *Liometopum* bodies, versus *Liometopum* guts and *Platyusa* bodies | 228 *Liometopum* guts vs. *Liometopum* bodies distances, 60 *Liometopum* guts vs. *Platyusa* bodies distances (derived from 12 *Liometopum* guts, 19 *Liometopum* bodies, and 5 *Platyusa* bodies) | | $2.5125 \times 10^{-6}$ | W = 3935 |
| | Weighted UniFrac distances between *Liometopum* guts and *Liometopum*, versus *Liometopum* guts and *Platyusa* guts | 228 *Liometopum* guts vs. *Liometopum* bodies distances, 60 *Liometopum* guts vs. *Platyusa* guts distances (derived from 12 *Liometopum* guts, 19 *Liometopum* bodies, and 5 *Platyusa* guts) | | $6.9978 \times 10^{-6}$ | W = 4129 |
| **Figure S5** | **Comparison** | **N number** | **Methods** | ***P* value** | **Statistical values** |
| S5 | Weighted UniFrac distances between *Liometopum* and *Sceptobius*, versus *Platyusa* and *Pella* | 868 *Liometopum* vs. *Sceptobius* distances, 26 *Pella* vs. *Platyusa* distances (derived from 31 *Liometopum* samples, 28 *Sceptobius* samples, 2 *Pella* samples, and 13 *Platyusa* samples) | Wilcoxon test with the Benjamini-Hochberg method for controlling the false discovery rate | 0.7534 | W = 10876 |

| | | | | | |
|---|---|---|---|---|---|
| | Weighted UniFrac distances between *Platyusa* and *Pella*, versus *Lissagria* and *Sceptobius* | 26 *Pella* vs. *Platyusa* distances, 56 *Lissagria* vs. *Sceptobius* distances (derived from 2 *Pella* samples, 13 *Platyusa* samples, 2 *Lissagria* samples, and 28 *Sceptobius* samples) | | 0.4194 | W = 619 |
| | Weighted UniFrac distances between *Lissagria* and *Sceptobius*, versus *Platyusa* and *Sceptobius* | 56 *Lissagria* vs. *Sceptobius* distances, 364 *Platyusa* vs. *Sceptobius* distances (derived from 2 *Lissagria* samples, 28 *Sceptobius* samples, and 13 *Platyusa* samples) | | 0.2518 | W = 8730 |
| | Weighted UniFrac distances between *Platyusa* and *Sceptobius*, versus *Liometopum* and *Lissagria* | 364 *Platyusa* vs. *Sceptobius* distances, 62 *Liometopum* vs. *Lissagria* distances (derived from 28 *Sceptobius* samples, 13 *Platyusa* samples, 31 *Liometopum* samples, and 2 *Lissagria* samples) | | 0.4745 | W = 10522 |
| | Weighted UniFrac distances between *Liometopum* and *Lissagria*, versus *Liometopum* and *Platyusa* | 62 *Liometopum* vs. *Lissagria* distances, 403 *Liometopum* vs. *Platyusa* distances (derived from 31 *Liometopum* samples, 2 *Lissagria* samples, and 13 *Platyusa* samples) | | 0.2957 | W = 11067 |
| | Weighted UniFrac distances between *Liometopum* and *Platyusa*, versus *Lasius* and *Pella* | 403 *Liometopum* vs. *Platyusa* distances, 10 *Lasius* vs. *Pella* distances (derived from 31 *Liometopum* samples, 13 *Platyusa* samples, 5 *Lasius* samples, and 2 *Pella* samples) | | $8.0199 \times 10^{-5}$ | W = 391 |
